## Supplementary File Guide for "Engineered transcription-associated Cas9 targeting in eukaryotic cells"

**Title: Supplementary Information**
**Description:** Combined PDF containing Supplementary Figs. 1–16, Supplementary Notes 1–6, and Supplementary References. Design considerations for engineering TraCT are outlined in Supplementary Notes 1–2; additional experimental considerations and caveats are elaborated in Supplementary Notes 3–6.

**Title: Supplementary Data 1**
**Description:** MNase-seq analysis of β-E-induced changes at individual nucleosomes within the *LYS2* ORF and across chromosome II

**Title: Supplementary Data 2**
**Description:** List of *S. cerevisiae yeast* strains used in this work

**Title: Supplementary Data 3**
**Description:** Circular plasmid DNA used in this work

**Title: Supplementary Data 4**
**Description:** Sequences of commercially synthesized linear single-stranded DNA oligonucleotides used in experimental assays and human cell culture work

**Title: Supplementary Data 5**
**Description:** Linear double-stranded DNA fragments used for CRISPR-assisted strain construction or donor-dependent editing assays in yeast

**Title: Supplementary Data 6**
**Description:** Raw results from analyses of human cell culture deep sequencing data (output from IDT's CRISPR Analysis Tool).
