## Supplementary Information for "Engineered transcription-associated Cas9 targeting in eukaryotic cells"

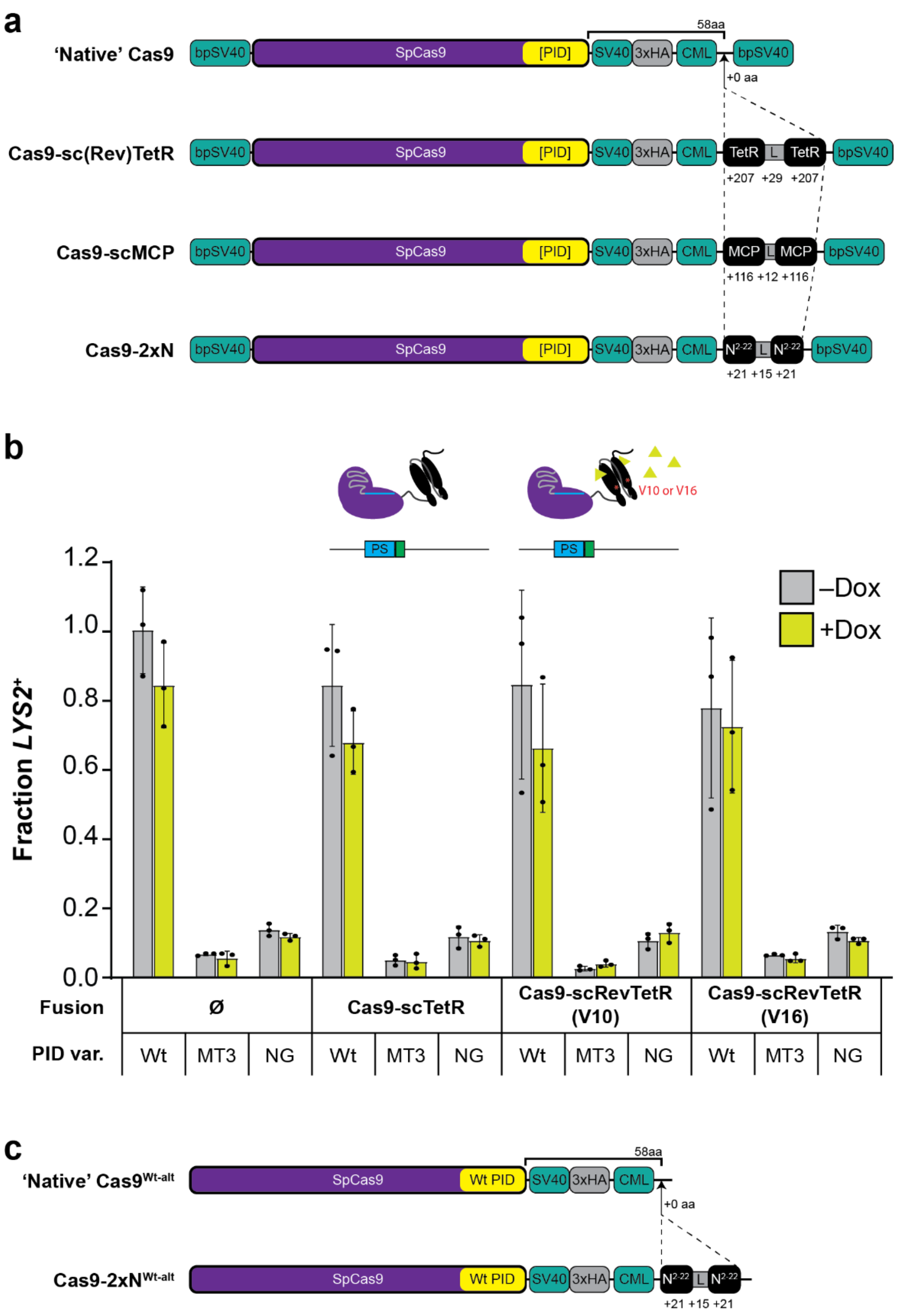


**Supplementary Fig. 1 | Summary of Cas9 fusion proteins used in this work and control experiments for sc(Rev)TetR performed in a strain lacking *tetO2*. a**, One-dimensional protein schematics diagramming the native (d)Cas9 scaffold and isogenic insertion site for (d)Cas9 fusions tested in yeast. A previously described[42] 58 amino acid (aa) linker that includes SV40 and C-myc-like (CML) nuclear localization signals (NLSs) is fused to *Streptococcus pyogenes* Cas9 (SpCas9) immediately downstream of its C-terminal PAM-interacting domain (PID). The additional aa lengths of each fusion domain segment are indicated below; e.g., +29 for the linker (L) connecting tandem TetR monomers. Additional bipartite SV40 (bpSV40) NLSs are present at the extreme N- and C-termini of all Cas9 proteins. 3xHA; three tandem repeats of the influenza hemagglutinin tag. **b**, Transformation-associated editing with the Cas9-sc(Rev)TetR fusions or native Cas9 control (Ø) in yeast lacking a *tetO2* element. Measurements were performed as in Fig. 1b. Inset schematics depict binding-proficient TetR (–Dox) and RevTetR (+Dox) states that remain unbound in the absence of a *tetO2* site downstream of the protospacer (PS). Error bars, mean ± s.d. ($n$ = 3, biological replicates). **c**, One-dimensional protein schematics with labeling as in panel **a** but for the alternative Cas9 architectures expressed in human cells without bpSV40 NLSs; only proteins with Wt PIDs were constructed.

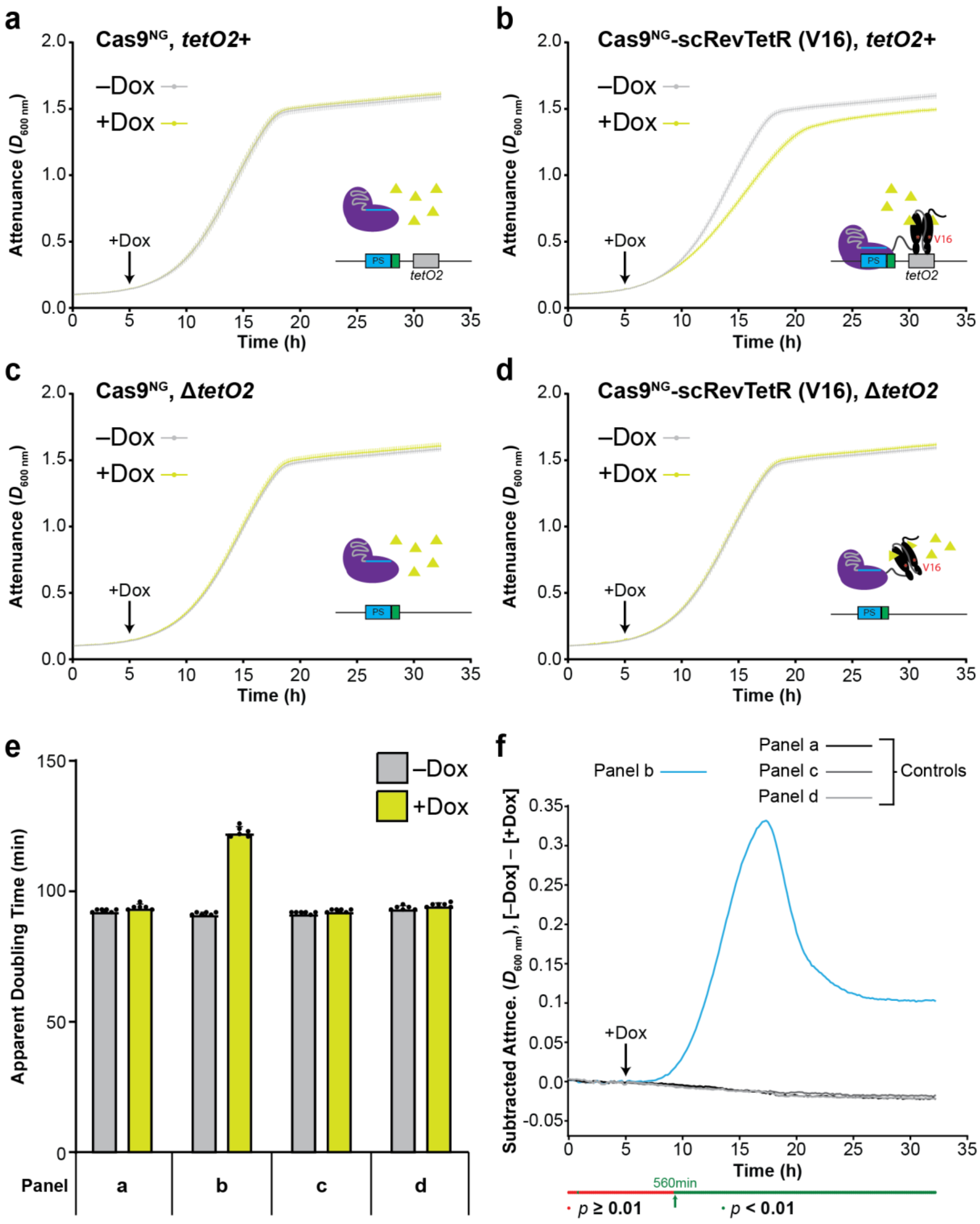


**Supplementary Fig. 2 | Time-resolved detection of a Dox-induced targeting phenotype for Cas9$^{NG}$-scRevTetR (V16) in liquid culture assays. a**, Growth of the specified strain in liquid cultures with (+Dox) or without (–Dox) induction at the indicated 5 h timepoint (black arrow) was monitored at high resolution using a plate reader. Inset illustration depicts a native Cas9$^{NG}$ protein that is unable to bind the *tetO2* element in the presence of Dox because it lacks the scRevTetR fusion. Attenuance ($D_{600\ nm}$)

readings were taken every 10 min. Error bars, mean ± s.d. ($n = 6$, biological replicates). **b**, Same experimental setup as in **a** except the strain tested contains a scRevTetR (V16) fusion that is bound in the presence of Dox, as shown in the inset. **c**, Same experimental setup as in **a** except the strain tested lacks the *tetO2* element. **d**, Same experimental setup as in **b** except the strain tested lacks the *tetO2* element and thus its scRevTetR domain remains unbound even in the presence of Dox. **e**, Apparent doubling times calculated from all growth curves plotted above (panels **a**-**d**). **f**, Time-resolved detection of Dox-induced growth phenotypes. Differences in growth were quantified at each time point for panels **a-d** above by subtracting the $D_{600\ nm}$ readings of +Dox cultures from those of –Dox cultures. Significant differences calculated for panel **b** ($p < 0.01$) at each time point are denoted with green dots below the plot and were first detected at 560 min growth. To improve readability, only the mean differences calculated for each panel are plotted.

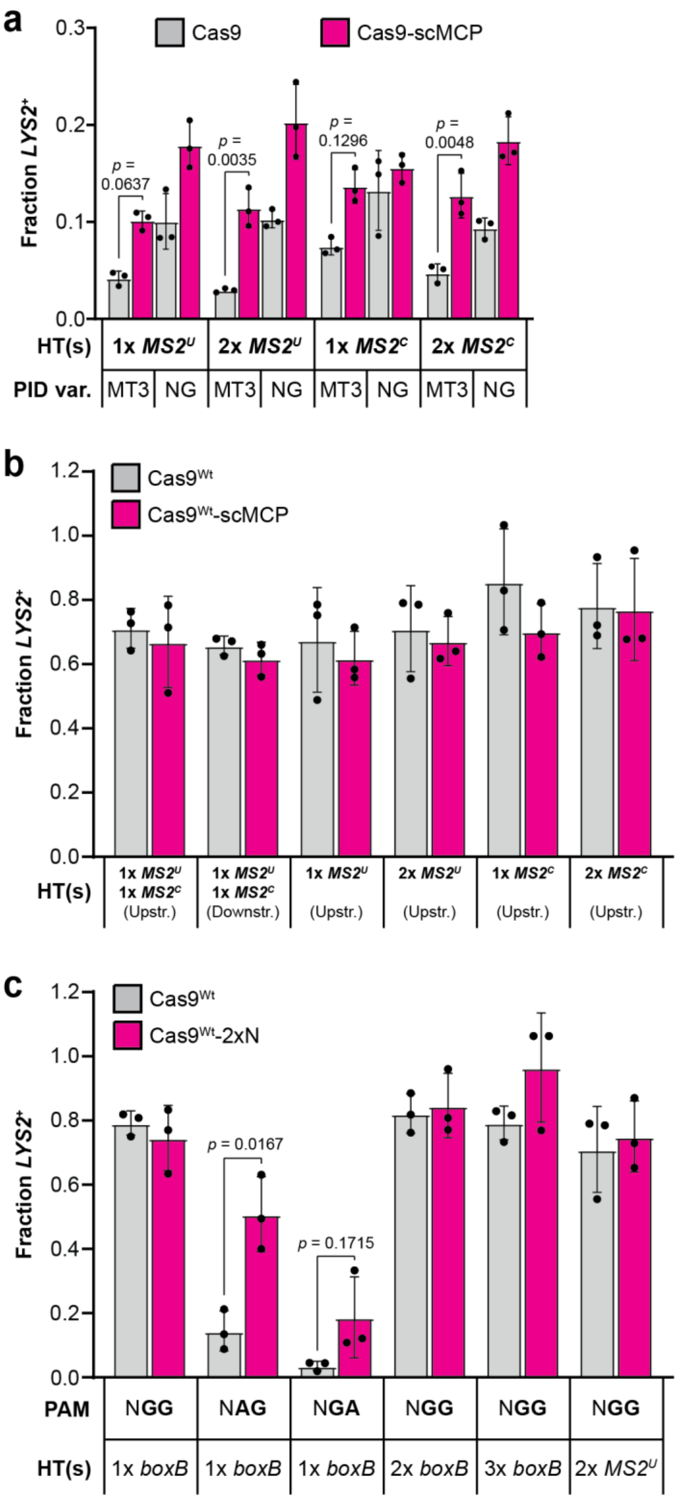


**Supplementary Fig. 3 | Testing of alternative MS2 hairpin configurations, Wt PID controls, and Cas9$^{Wt}$-2xN with non-NGG subPAMs. a**, Transformation-associated editing fractions measured for MT3 and NG variants of Cas9-scMCP or native Cas9 in strains containing alternative HT configurations as indicated. Error bars, mean ± s.d. (*n* = 3, biological replicates). **b**, Control experiments with Wt PIDs, using the same setup and HT strains tested in panel **a** and Fig. 1e. Error bars, mean ± s.d. (*n* = 3, biological replicates). **c**, Experiments with Wt PID variants of Cas9-2xN or native Cas9 using the same setup and HT strains tested in Fig. 1f, as well as two additional strains containing one *boxB* HT and either NAG or NGA PAMs as indicated. Error bars, mean ± s.d. (*n* = 3, biological replicates).

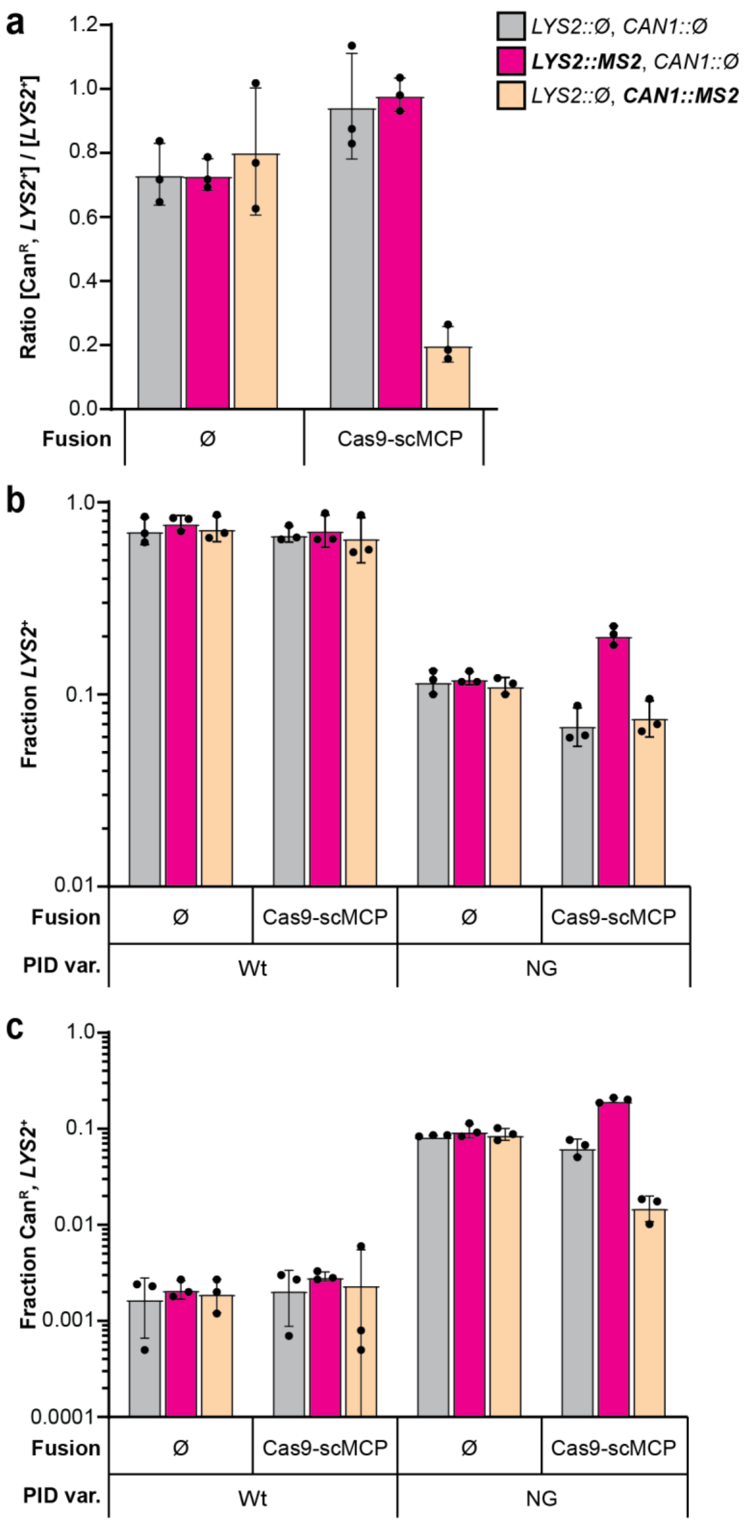


**Supplementary Fig. 4 | Re-plotting of data from Fig. 2 and testing of Wt PID controls in dual-reporter strains. a**, Raw counts of *LYS2*$^{+}$ and Can$^{R}$, *LYS2*$^{+}$ colonies from the experiments of Figs. 2b–c were used to calculate and plot ratios as indicated (Can$^{R}$, *LYS2*$^{+}$ counts over *LYS2*$^{+}$ counts). Error bars, mean ± s.d. (*n* = 3, biological replicates). **b**, Re-plotting of data from Fig. 2b in logarithmic scale, alongside data from Wt PID control experiments. Error bars, mean ± s.d. ($n$ = 3, biological replicates). **c**, Re-plotting of data from Fig. 2c in logarithmic scale, alongside data from Wt PID control experiments. Error bars, mean ± s.d. ($n$ = 3, biological replicates).

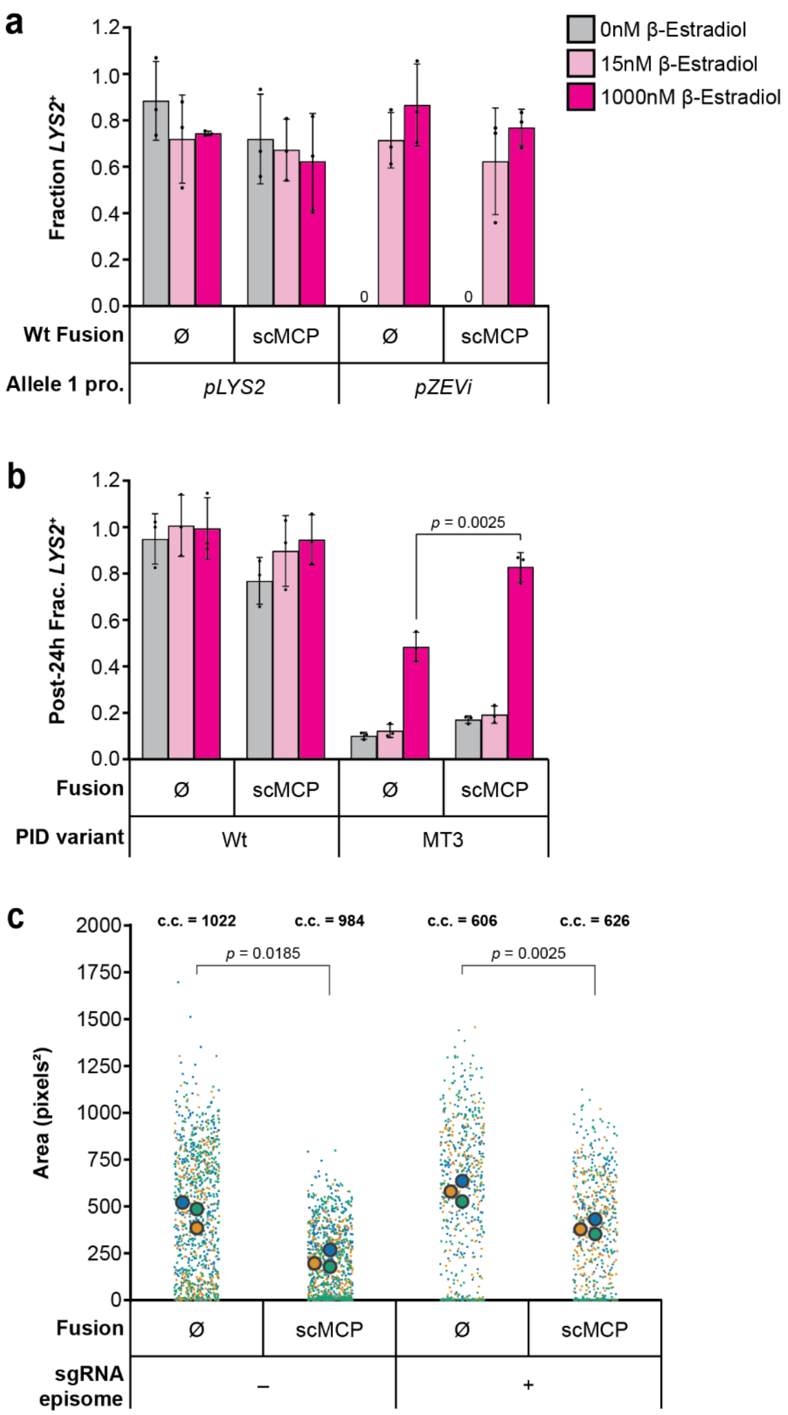


**Supplementary Fig. 5 | Control experiments for Fig. 3 and colony size comparisons for strains expressing native Cas9$^{Wt}$ or Cas9$^{Wt}$-scMCP under non-targeting conditions. a**, Transformation-associated editing fractions measured under the same conditions as Fig. 3b experiments, but with Wt PID variants of Cas9-scMCP or native Cas9 in *pLYS2* or *pZEVi* haploid strains. Zero (0) *LYS2*$^{+}$ colonies of the *pZEVi* strain were detected in the absence of β-E at the time of scoring. Error bars, mean ± s.d. ($n$ = 3, biological replicates). **b**, Diploid *LYS2*$^{+}$ fractions assayed with β-E at each concentration (color coded as in panel **a**) by plating the Cas9$^{MT3}$ and Cas9$^{MT3}$-scMCP transformant populations of Fig. 3c after the 24 h liquid outgrowth step and immediately before the subculture induction step. Wt PID control populations were also assayed here post-outgrowth, though not subcultured or studied further. All *LYS2*$^{+}$ colonies detected in the absence of β-E are inferred to result from repair of the constitutive *pLYS2* allele since repair of the *pZEVi* allele in non-induced haploids is insufficient to generate *LYS2*$^{+}$ colonies under the conditions tested. Error bars, mean ± s.d. ($n$ = 3, biological replicates). **c**, Colony sizes measured for BY4741 strains carrying the indicated Cas9 plasmids with (+) or without (–) episomal expression of an sgRNA, which, is effectively non-targeting because this background lacks the protospacer insertion in *LYS2*. The total number of colonies counted (c.c.) for each strain is indicated in boldface above each column. Datapoints from each biological replicate ($n$ = 3) are color-coded, and the corresponding mean values measured for each replicate are indicated with large outlined dots. Parametric $t$ tests were performed on means (paired within replicates) to obtain the $p$ values shown.

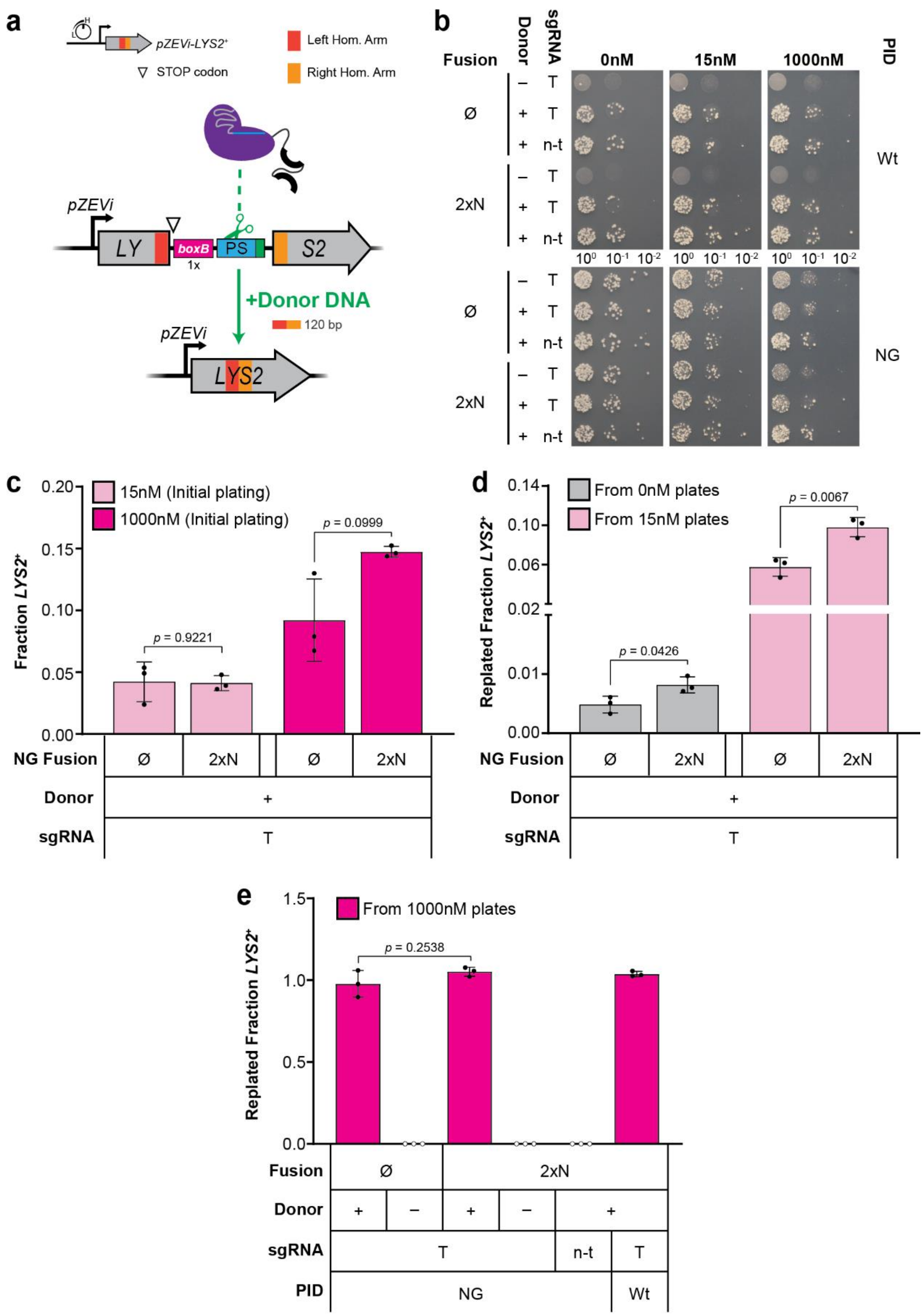


**Supplementary Fig. 6 | TraCT can stimulate donor-dependent editing in *pZEVi* haploid yeast strains. a**, Schematic summary of the donor-dependent *LYS2* editing system. Upon cleavage, the disrupted *pZEVi-LYS2* locus can be reconstituted through homology-directed repair with a 120 bp donor DNA fragment that scarlessly removes intervening sequences between the left (red) and right (orange) homology arms. **b**, Post-transformation colony phenotypes for strains harboring the Wt or NG PID variants of native Cas9 (Ø) or Cas9-2xN as indicated, plated on media with lysine to permit growth of all transformants. Transformations were performed with targeting (T) or non-targeting (n-t) sgRNA plasmids and co-delivered (+) or absent (–) donor DNA. The resulting populations were plated as serial dilutions on plates that contained β-E at the indicated concentrations (0nM, 15nM or 1000nM) in addition to lysine. Photos shown contain representative data from one of the three biological replicates ($n$ = 3). **c**, Transformation-associated editing fractions measured for the indicated NG populations from panel **b**; plating was performed in parallel from the same serial dilutions but using lysine-free media with β-E at the indicated concentrations. Error bars, mean ± s.d. ($n$ = 3, biological replicates). **d**, *LYS2*$^{+}$ fractions measured for the indicated NG populations from panel **b**, but after an outgrowth step with lysine present on plates containing β-E at the indicated concentrations. To activate *LYS2* expression in edited cells during the replating step, lysine-free plates contained 1000nM β-E. **e**, *LYS2*$^{+}$ fractions measured for the indicated NG populations from panel **b** as performed in panel **d**, except that 1000nM β-E was used during the outgrowth step.

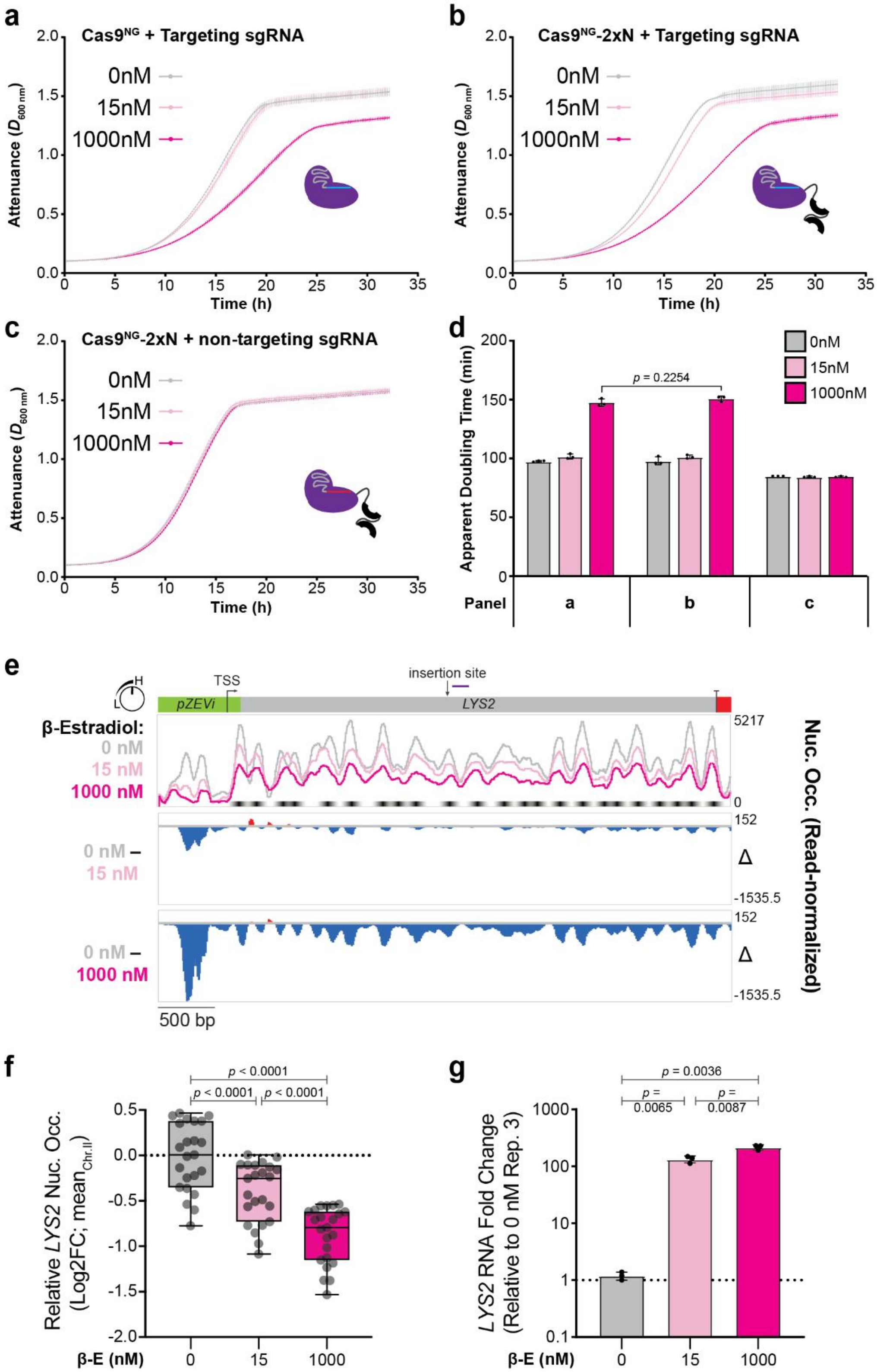


**Supplementary Fig. 7 | Transcriptional induction at the *LYS2* locus can increase basal Cas9$^{NG}$ targeting and reduce nucleosome occupancy. a**, Plate reader experiments to monitor growth in liquid media with β-E (15nM or 1000nM) or lacking it altogether (0nM). The *pZEVi* reporter strain tested here is an unedited isolate from donor-dependent editing experiments with native Cas9$^{NG}$; this protein lacks an auxiliary RBD fusion as shown in the inset illustration. Attenuance ($D_{600\ nm}$) readings were taken every 10 min. Error bars, mean ± s.d. ($n$ = 3, biological replicates). **b**, Same experimental setup as in **a** except the strain tested contains a 2xN fusion, as shown in the inset, that can interact with boxB RNA transcribed upstream of the protospacer. **c**, Same experimental setup as in **b** except the strain tested has non-targeting spacer sequences (red line) in its sgRNA scaffold. **d**, Apparent doubling times calculated from all growth curves plotted above (panels **a**-**c**). **e**, Nucleosome occupancies determined from MNase-seq experiments with 0nM, 15nM, or 1000nM β-E induction were calculated after normalizing across samples for read depth and then plotted for the intact *LYS2* ORF (gray), *pZEVi* promoter (green), and native *LYS2* terminator (red) in scale with the schematic above. To emphasize changes in read-normalized nucleosome occupancies at each β-E concentration, the lower two panels plot differences (Δ) relative to 0nM as indicated. Bent arrow in *pZEVi* denotes position of the transcriptional start site (TSS), and straight arrow above *LYS2* denotes the site where insertions begin in target strains. The inferred positions of 23 nucleosomes within the *LYS2* ORF are indicated with black spots below the top panel coverage traces. Relative percentages of read-normalized nucleosome occupancies were also calculated (15nM / 0nM or 1000nM / 0nM) for each of the 23 nucleosomes (see Supplementary Data 1). All data plotted represent means calculated from $n$ = 3 biological replicates. Purple line above *LYS2* and downstream of the target insertion site delineates the amplicon generated in qRT-PCR (see panel **g**). **f**, Box and whisker plot of read-normalized occupancies for *LYS2* ORF nucleosomes are presented for each β-E concentration in terms of a Log2 Fold Change (Log2FC). Log2FC values were calculated relative to the mean read-normalized nucleosome occupancy measured across chromosome II (chr.II) at each concentration (denoted with a black dotted line). Gray dots represent mean Log2FC values calculated for each of the 23 nucleosomes from $n$ = 3 biological replicates. Significant differences in Log2FC across the entire *LYS2* ORF at each β-E concentration were determined with a Tukey's test. **g**, qRT-PCR experiments to determine the fold change in *LYS2* RNA levels for MNase-seq cultures at each β-E concentration. Fold changes were calculated relative to the lowest of three values measured at 0nM β-E, which was normalized to '1' (black dotted line). Considering that the maximal RNA levels previously obtained with a *pZEVi* promoter were ~ 100-fold increased after 2 h induction[65], *LYS2* transcript accumulation is likely already saturated at our timepoint. Error bars, mean ± s.d. ($n$ = 3, biological replicates).

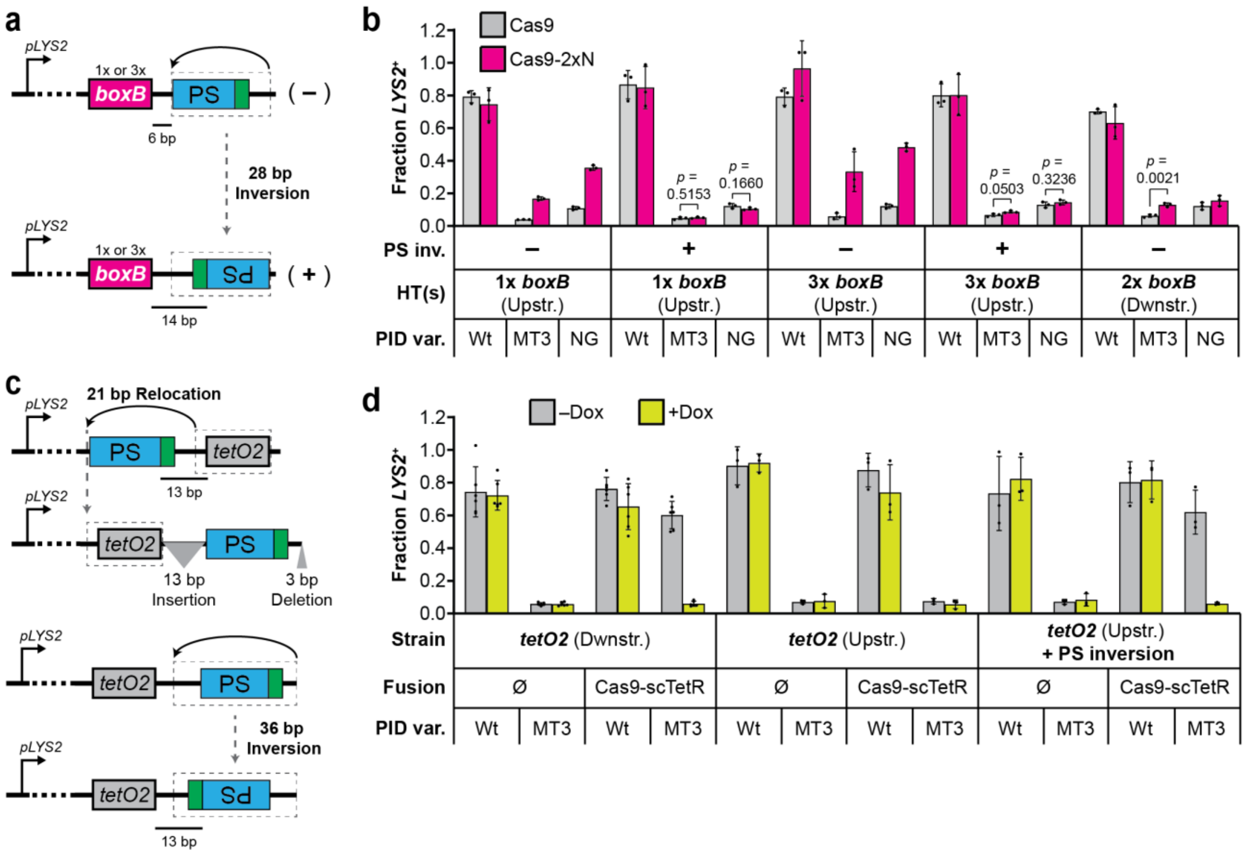


**Supplementary Fig. 8 | Protospacer orientation can influence TraCT activity. a**, Schematic summary of the 28 bp target inversion (dashed gray box) that includes the protospacer and 8 adjacent bp on its PAM side, including the NGG PAM, in SSA reporter strains with *pLYS2* and 1x or 3x *boxB* HTs upstream of the target. Distance in bp between the protospacer and nearest upstream *boxB* HT is also shown. **b**, Transformation-associated editing fractions measured for the indicated PID variants of native Cas9 or Cas9-2xN in strains with the inverted (+) or parental (–) protospacer orientation and various HT configurations. Error bars, mean ± s.d. ($n$ = 3, biological replicates). **c**, Schematic summary of the *tetO2* relocation and protospacer inversion tested in SSA reporter strains with *pLYS2*. Top, 21 bp that included *tetO2* were relocated (dashed gray box) immediately upstream of the protospacer while 13 bp were inserted between *tetO2* and the protospacer to preserve spacing relative to the original *tetO2* (Dwnstr.) strain design; deletion of 3 bp at positions 9 through 11 downstream of the protospacer was intended to partially offset the SSA reporter insertion's overall increase in length. Bottom, 36 bp that included 8 bp on each side of the protospacer were inverted (dashed gray box) relative to the relocated *tetO2* (Upstr.) design, thus preserving 13 bp of spacing between *tetO2* and the protospacer. **d**, Transformation-associated editing fractions measured for Wt and MT3 variants of native Cas9 (Ø) or Cas9-scTetR in panel **c** strains as indicated, with the same experimental setup used for Fig. 1b; Wt and MT3 data collected with the original *tetO2* (Dwnstr.) strain is replotted from Fib. 1b for comparison. Error bars, mean ± s.d. ($n$ = 6 or $n$ = 3, biological replicates).

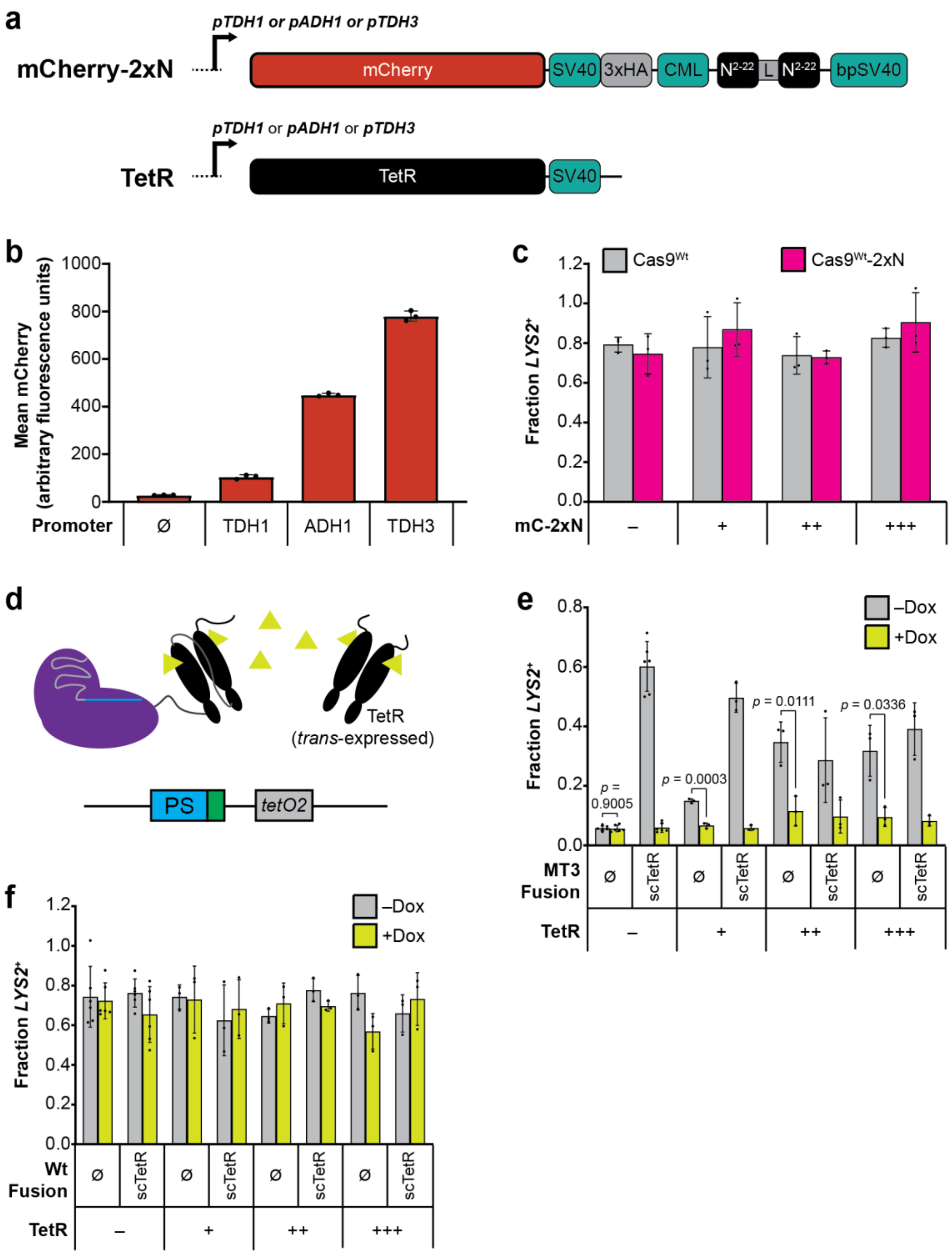


**Supplementary Fig. 9 | Additional summary and supporting experiments for Fig. 5, including tests with ectopically expressed TetR proteins in SSA reporter strains. a**, One-dimensional protein schematics diagramming features of the nuclear-localized mCherry-2xN (mC-2xN) and TetR proteins expressed from the ectopic *YKL162C-A* locus on chromosome XI, using one of three promoters as indicated above the bent arrow. C-terminal protein sequences beyond mCherry in mC-2xN are identical to sequences beyond Cas9-2xN's PID and include the 58 aa linker; the same abbreviations from Supplementary Fig. 1 are used here. **b**, mCherry fluorescence measured for Fig. 4b yeast strains expressing mC-2xN from each of the indicated promoters; background fluorescence measured with the isogenic strain lacking an mC-2xN expression cassette is also plotted (Ø). Mean values were calculated from *n* = 3 (biological replicates) populations of single yeast cells cultured in parallel and analyzed by flow cytometry on the same day. Error bars, mean ± s.d. **c**, Transformation-associated editing fractions measured with the same strains and conditions tested in Fig. 4b but after delivery of $Cas9^{Wt}$ or $Cas9^{Wt}$-2xN. For comparison, data shown for the strain lacking mC-2xN (–) are replotted from Supplementary Fig. 3c (NGG, 1x *boxB*). Error bars, mean ± s.d. (*n* = 3, biological replicates). **d**, Schematic illustration summarizing the expected effect of Dox induction in strains containing $Cas9^{MT3}$-scTetR, *trans*-expressed TetR, and a protospacer-proximal *tetO2* element. **e**, Transformation-associated editing with native $Cas9^{MT3}$ or the $Cas9^{MT3}$-scTetR fusion in strains with a protospacer-proximal *tetO2* element and TetR absent (–) or expressed in *trans* at the indicated levels (+/++/+++). Measurements were performed as in Fig. 1b; MT3 data with strains lacking *trans*-expressed TetR are replotted from there for comparison. Error bars, mean ± s.d. (*n* = 6 or *n* = 3, biological replicates). **f**, Same experimental setup and strains tested in panel **e** but with $Cas9^{Wt}$ or $Cas9^{Wt}$-scTetR; the strain lacking *trans*-expressed TetR (–) is replotted from Wt data of Fig. 1b for comparison.

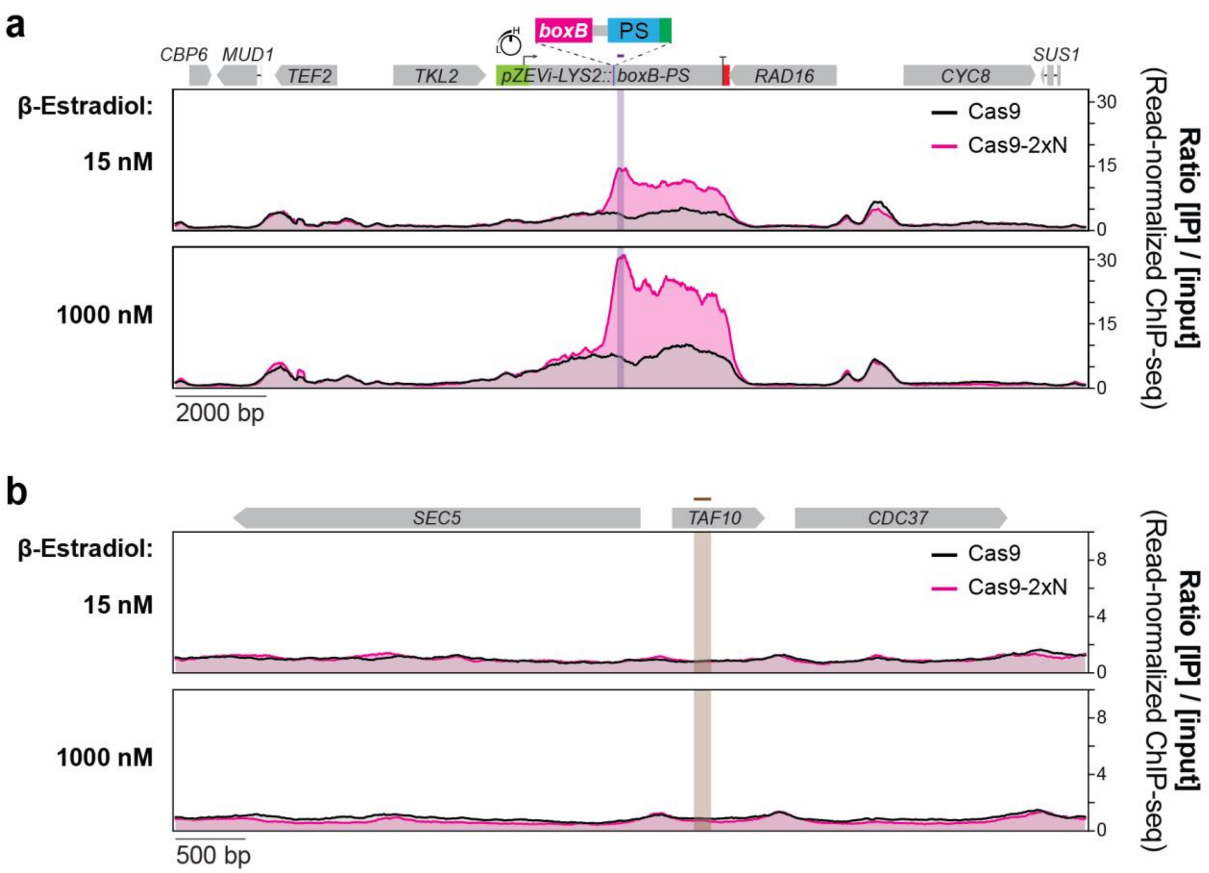


**Supplementary Fig. 10 | ChIP-seq coverage plots for Cas9$^{NG}$ and Cas9$^{NG}$-2xN that include control regions outside the *boxB*-tagged *pZEVi-LYS2* locus. a**, ChIP-seq data from the 15 and 1000nM β-E samples analyzed in Fig. 5c (*n* = 1, biological replicates), re-analyzed to plot coverage ratios across a larger region of chromosome II that includes flanking genes. Color coding is presented as in Fig. 5c, with grey elements in the scaled linear map atop the plots used to indicate all the full-length non-dubious coding sequences of this region. The non-targeting sgRNA used in these experiments prevents cleavage at the protospacer (PS). **b**, ChIP-seq data from the same samples analyzed in panel **a** (*n* = 1, biological replicates) but re-analyzed to plot coverage ratios across a region of chromosome IV that includes the *TAF10* gene. The 121 bp region probed by qPCR in Fig. 5b is delineated with brown shading and a brown line above *TAF10* in the scaled linear map atop the plots.

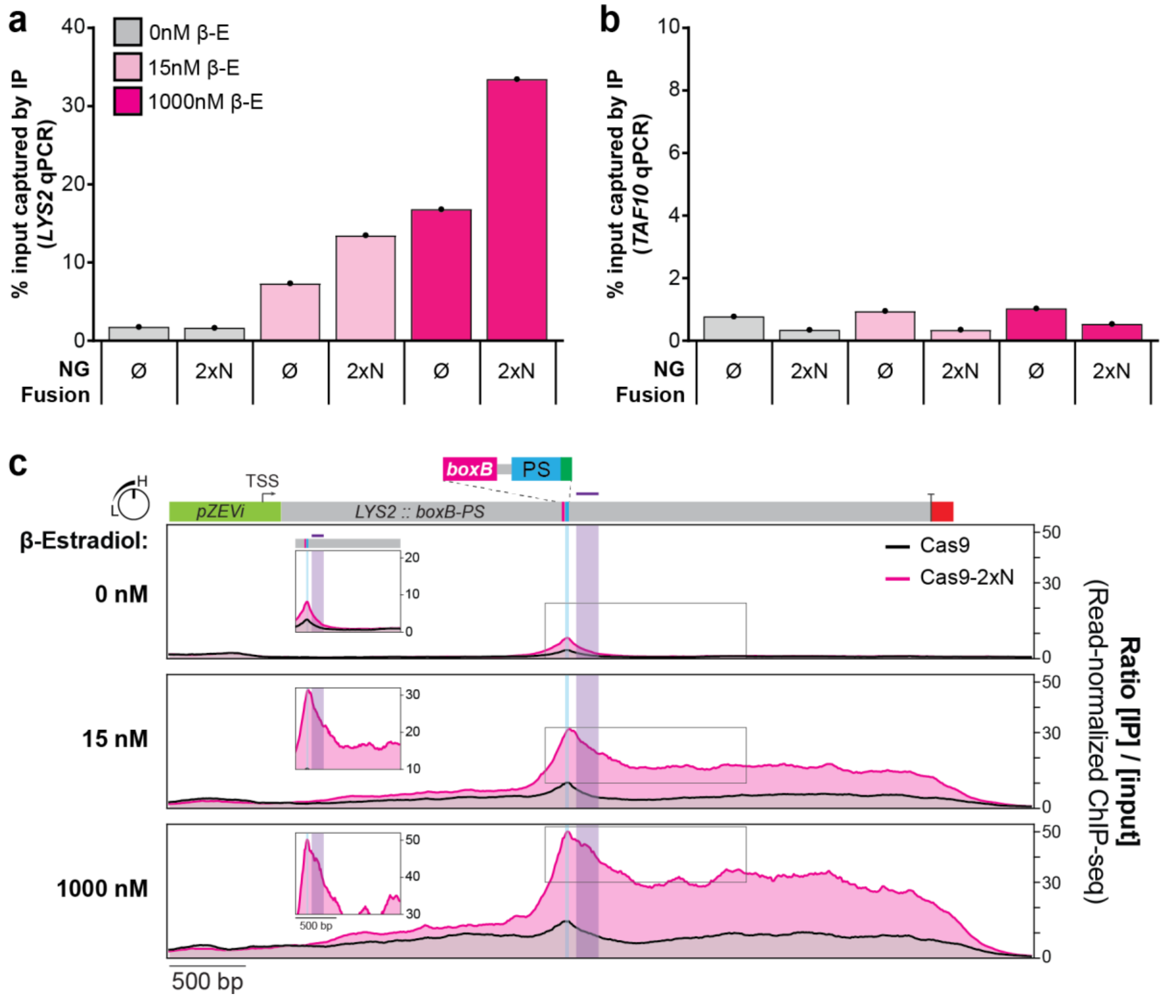


**Supplementary Fig. 11 | Accumulation on or near protospacer DNA is higher for dCas9-2xN than for native dCas9 at transcribed targets with an upstream *boxB*. a**, ChIP-qPCR for yeast strains carrying a targeting sgRNA and dCas9$^{NG}$ or dCas9$^{NG}$-2xN (both of which bear a 3xHA tag, see Supplementary Note 1), either untreated (0nM) or treated (15 and 1000nM) with β-E for transcriptional induction of the *boxB*-tagged *pZEVi-LYS2* locus. The percentage of chromatin captured by immunoprecipitation (IP) with anti-HA antibody out of total input chromatin was quantified using *LYS2*-specific primers for an amplicon within 250 bp downstream of *boxB*. Data were collected from single experiments (*n* = 1, biological replicates). **b**, Background control reactions performed in parallel for all panel **a** samples, with qPCR primers amplifying the *TAF10* locus on a separate chromosome that lacks *boxB*. Data were collected from single experiments (*n* = 1, biological replicates). **c**, ChIP-seq enrichments across the *boxB*-tagged *pZEVi-LYS2* locus, determined with anti-HA antibody for dCas9$^{NG}$ and dCas9$^{NG}$-2xN at each β-E concentration. Data were generated from the samples tested in panel **a** and plotted as the ratio of IP coverage over input coverage after normalizing each sample for read depth (*n* = 1, biological replicates). The region downstream of *boxB* that was probed by qPCR is delineated with purple shading and a purple line above *LYS2* in the scaled linear map atop the plots. Blue shading delineates the 20 bp protospacer region. Calculating IP/input coverage ratios at every single bp across the protospacer revealed 2.69-, 3.02-, and 3.34-fold higher mean pull-down for dCas9$^{NG}$-2xN relative to dCas9$^{NG}$ with 0, 15, and 1000nM β-E, respectively. Boxed insets are rescaled to dimensions that emphasize relative differences between the maximum values at the protospacer and other local maxima further downstream. TSS, transcriptional start site; PS, protospacer

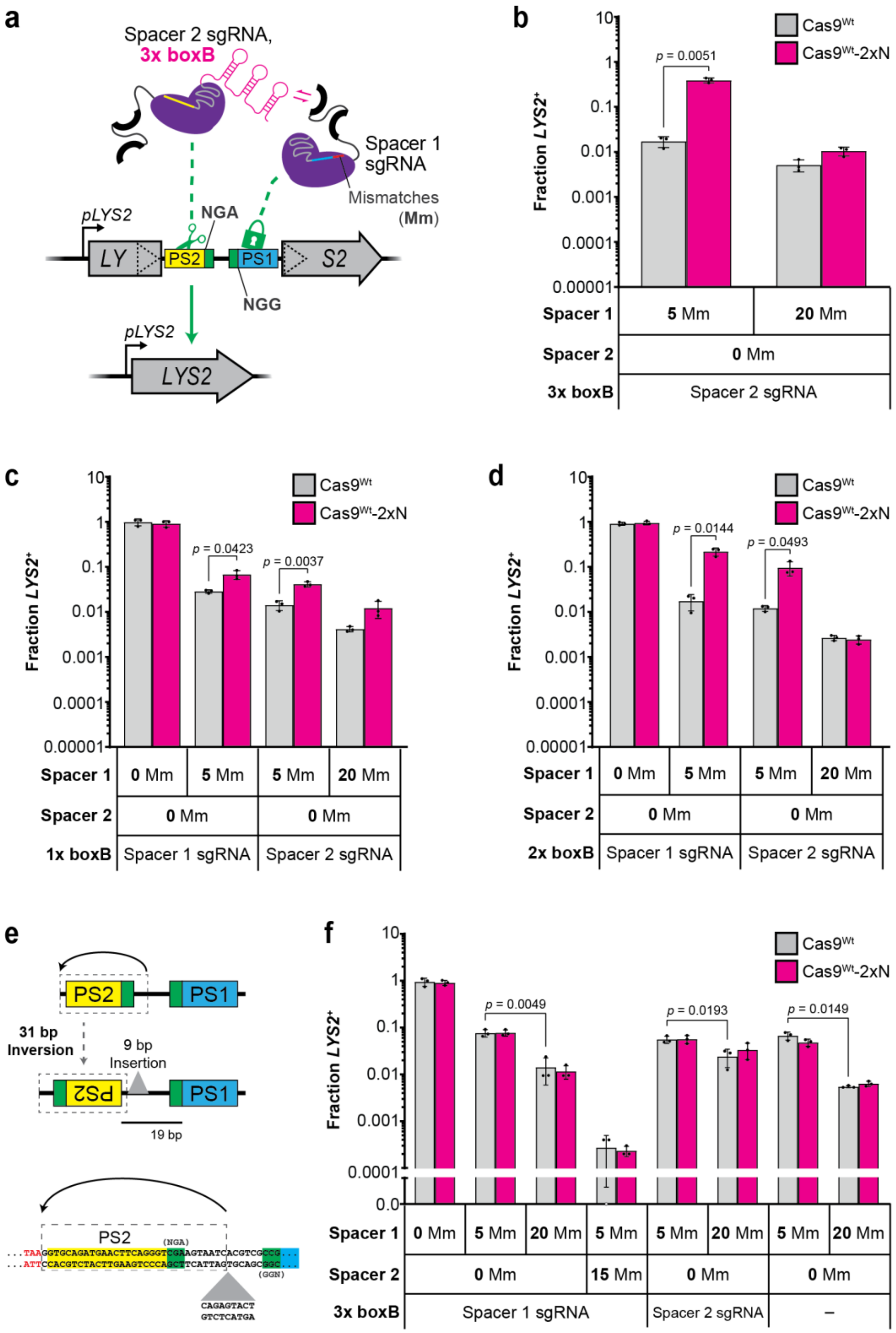


**Supplementary Fig. 12 | Testing of alternative sgRNA configurations and target orientations with the paired-protospacer system for recruitment via anchored boxB hairpins. a**, Schematic summary of the same SSA reporter strain used in Fig. 6 but tested with an alternative sgRNA configuration where Spacer 2 is fused to the 3x boxB scaffold. **b**, Transformation-associated editing fractions measured for Cas9$^{Wt}$-2xN or native Cas9$^{Wt}$ with the sgRNA configuration of panel **a** and 5 or 20 Mm in Spacer 1 as indicated. Error bars, mean ± s.d. ($n$ = 3, biological replicates). **c**, Transformation-associated editing fractions measured as in panel **b** but with a single boxB hairpin fused to the indicated sgRNA scaffold and spacer configurations. **d**, Same experimental setup as in panel **c** but with two boxB hairpins fused to sgRNA scaffolds. **e**, Schematic summary of the 31 bp target inversion (dashed gray box) that includes Protospacer 2 (PS2) and 10 adjacent bp on its PAM side, including the NGA PAM. A 9 bp insertion (gray triangle) was also introduced to preserve the 19 bp spacing between PS2 and Protospacer 1 (PS1). Sequence of the original strain is shown 5' to 3' at the bottom of the schematic, beginning with the upstream TAA stop codon (red text) and ending with the NGG PAM of PS1 (blue highlight). **f**, Same plasmids and experimental setup tested in panel **b** and Fig. 6b but with the PS2-inverted strain as summarized in panel **e**.

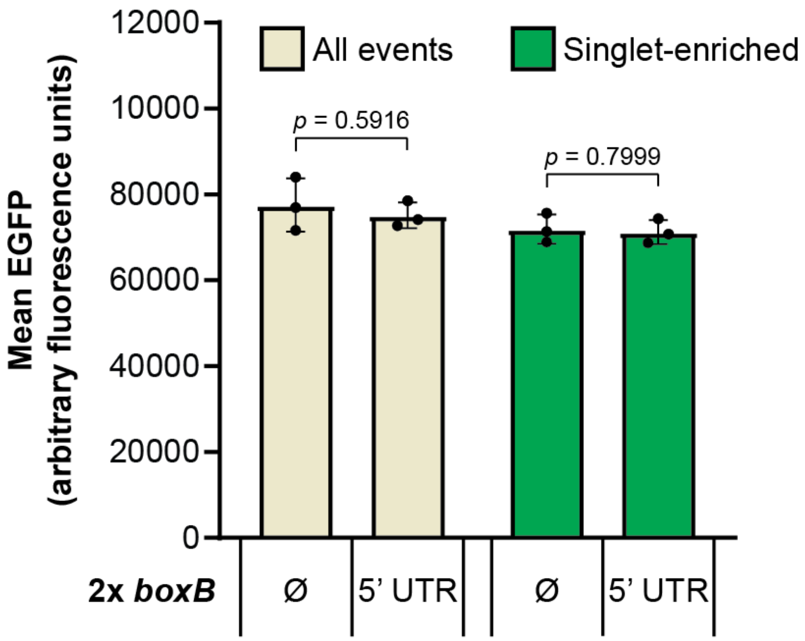


**Supplementary Fig. 13 | Baseline EGFP fluorescence was unaltered by insertion of *boxB*$^{2x}$ in the U2OS.EGFP::5’ UTR-*boxB*$^{2x}$ cell line.** Comparison of EGFP fluorescence levels in the human U2OS.EGFP parent line lacking *boxB* (Ø) and its derivative with two *boxB* HTs inserted upstream of the *EGFP* ORF (5’ UTR), measured under the same growth conditions used for indel assays. Mean values were calculated from $n$ = 3 (biological replicates) populations of ungated (All events) or gated (Singlet-enriched) cells that were cultured in parallel and analyzed by flow cytometry on the same day. Error bars, mean ± s.d.

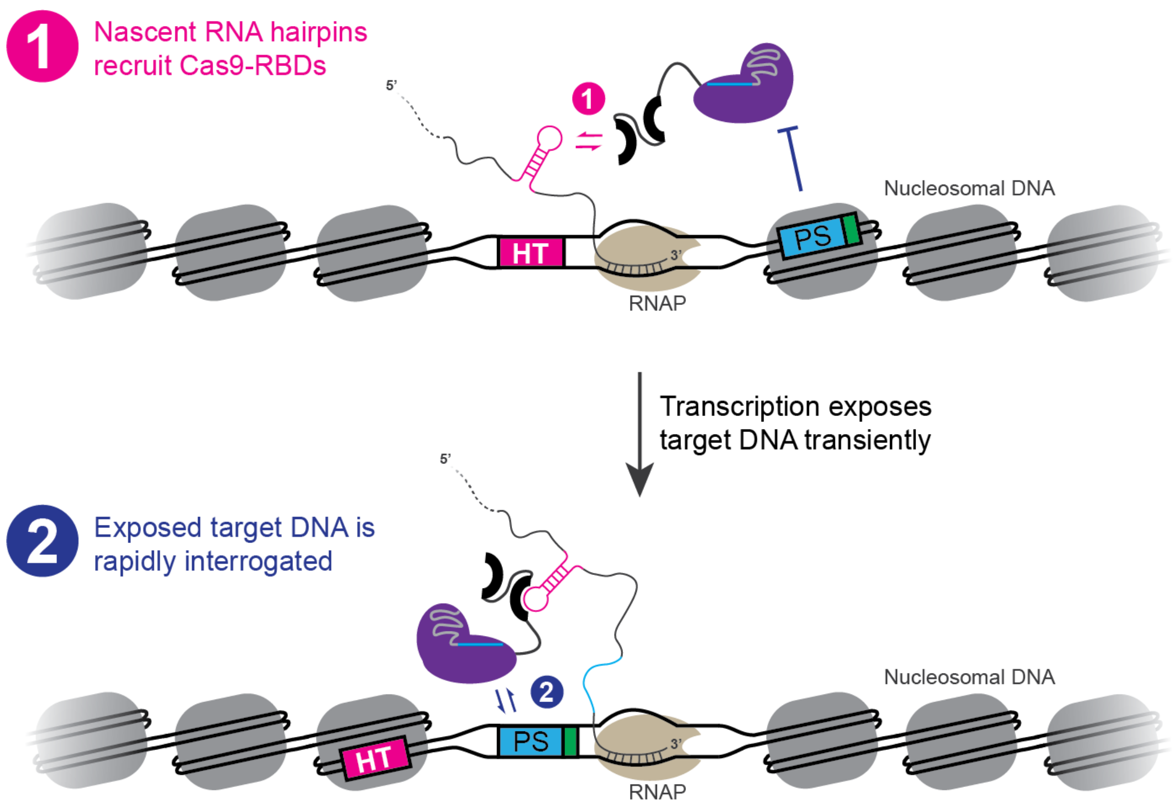


**Supplementary Fig. 14 | Schematic illustration of our proposed two-part model for engineered TraCT.** Top, nascent RNA hairpins are transcribed from HTs and recruit Cas9-RBDs upstream of a protospacer (PS) that is initially inaccessible due to occlusion of its subPAM (green) within nucleosomal DNA. Bottom, subsequent transcription by the RNA polymerase (RNAP) displaces downstream nucleosomes to transiently expose the subPAM and PS DNA. Tethering via the nascent RNA allows Cas9-RBDs to rapidly interrogate the target DNA while it is exposed.

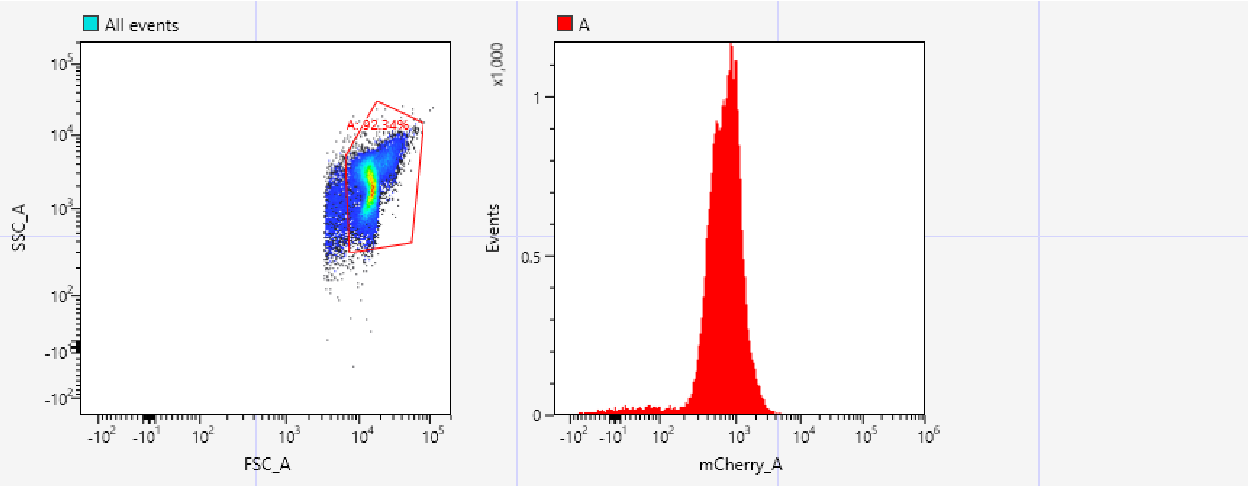


Gates and Statistics

| Name | Events | Parent% | Total% | Mean |
|---|---|---|---|---|
| All events | 32,490 | 100.00% | 100.00% | |
| A | 30,000 | 92.34% | 92.34% | |
| mCherry_A | | | | 776.95 |

**Supplementary Fig. 15 | Summary of the analytical gating strategy used in flow cytometry assays for measuring yeast fluorescence.** Top left, pseudo-colored density plot of all events detected, with gate ‘A’ capturing a majority of events in the expected range for forward scatter area (FSC_A) and side scatter area (SSC_A) measurements. Top right, histogram of mCherry-fluorescent events detected within the ‘A’ gate, determined from mCherry area (mCherry_A) measurements. Data shown are from an actual experiment with the *TDH3* promoter driving *mCherry-2xN* expression (Supplementary Fig. 9b); identical gates were applied whenever performing this assay.

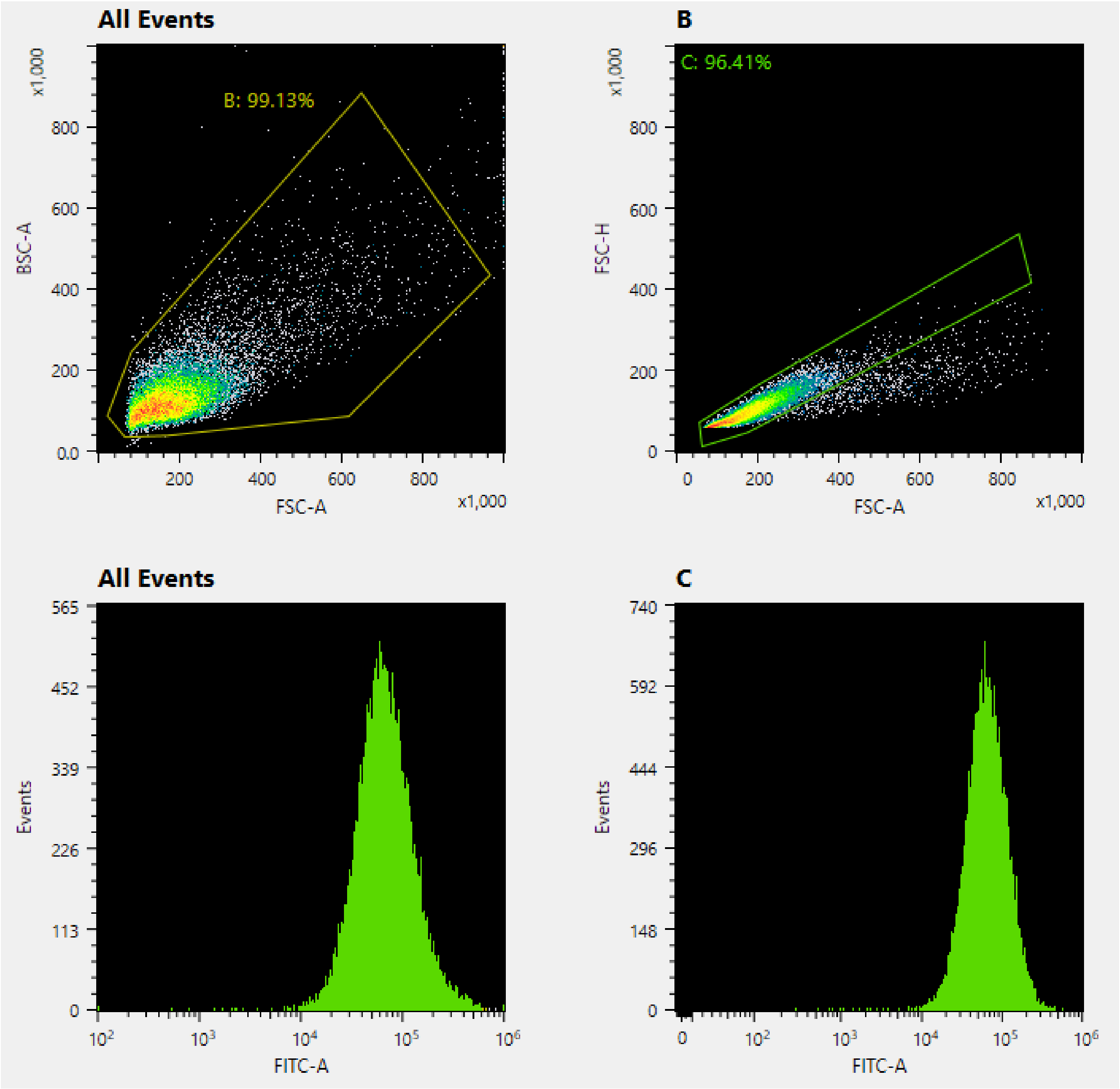


**Supplementary Fig. 16 | Summary of the analytical gating strategy used in flow cytometry assays for measuring human cell fluorescence.** Top, pseudo-colored density plots of all events (left) or 'B' events (right) gated from a majority population in the expected range for forward scatter area (FSC-A) and backscatter area (BSC-A) measurements. The 'C' gate was applied in the top right plot to enrich for singlet events, based on measurements of FSC-A and forward scatter height (FSC-H). Bottom, histogram of EGFP-fluorescent events detected within the ungated population (left) or within the 'C' gate (right), determined from FITC area (FITC-A) measurements that capture all emissions in the 500-550nm range. Data shown are from an actual experiment with the U2OS.EGFP parent line (Supplementary Fig. 13); identical gates were applied whenever performing this assay.

**Supplementary Note 1. Design of Cas9 fusion proteins generated in this work.**

All (d)Cas9 fusion proteins expressed in yeast were designed from a “native” (d)Cas9 scaffold essentially as encoded in the pNA0306 vector described previously[99], but with bipartite SV40 (bpSV40) nuclear localization signal (NLS) tags at each N- and C-terminus (Supplementary Fig. 1a). This scaffold also includes a previously described 58-residue linker[42] with SV40 and c-Myc-like NLS tags, as well as a 3xHA tag, inserted immediately downstream of the core SpCas9 open reading frame (ORF). All recombinant DNA or RNA binding domains were inserted immediately downstream of this 58-residue linker. Synonymous recoding of DNA encoding repeated protein sequences, where applicable, was used to minimize direct repeat homology tracts. The use of multiple NLS tags, including N- and C-terminal bpSV40 tags, was reported to enable robust nuclear localization of larger Cas9 fusion proteins[113].

The single-chain TetR (scTetR) and single-chain reverse TetR (scRevTetR) DNA binding domains were designed by fusing two TetR or two RevTetR monomers in tandem with a previously described 29-residue glycine/serine-rich linker[46] that comprises 6x consecutive $S_1G_4$ units when considering the C-terminal serine of the upstream monomer. N-terminal methionines were included with each monomer. The previously described V10 and V16 variants used for scRevTetR fusions are reviewed elsewhere[45].

The single-chain MS2 coat protein (scMCP) RNA binding domain was designed by fusing two MCP monomers in tandem with a 12-residue linker of the sequence A-(2x $S_1G_4$)-M. This linker length was chosen to exceed the length and presumed flexibility of linkers previously tested in tandem designs: A-M or A-L[114], G-A-P-G-I-H-P-G-M[115], A-

M[116]. Each MCP monomer included the V29I substitution that increases affinity for MS2 hairpins[117], as well as the dl-FG mutation reported to reduce or prevent higher-order multimerization required for capsid assembly without preventing dimerization and binding to MS2 hairpins[118,119].

The two tandem 21-residue $N^{2-22}$ peptides (2xN) for binding boxB RNA hairpins are derived from variants of bacteriophage lambda's N protein residues 2 through 22 with D2N,Q4R substitutions reported to improve affinity for boxB in vitro[63]. They were fused together with a 15-residue (3x)$G_4S_1$ linker as described previously in another work[85].

The Cas9-2xN$^{Wt\text{-}alt}$ fusion protein used in human cell culture experiments was derived from an alternative scaffold which served as its native Cas9$^{Wt\text{-}alt}$ control. Both lack bpSV40 NLSs but are otherwise similar to the Cas9 proteins expressed in yeast and contain the same 58-residue linker. The Cas9$^{Wt\text{-}alt}$ scaffold was derived from the Cas9$^{Wt}$-SV40-3xFLAG protein encoded in the previously described pGG439 plasmid[19] by replacing all C-terminally fused residues beyond SpCas9's core ORF with the 58-residue linker followed by 8 additional residues: T-E-R-P-L-R-G-S.

**Supplementary Note 2. Summary and DNA sequences of in vivo templates used for MS2 or boxB hairpin RNAs in this work.**

Our 2x *MS2* HT designs were based off sequences in the pIIIA/MS2 plasmids[120] which contained two *MS2*$^C$ HTs with a 20bp linker between them. Our *MS2*$^U$ HTs are in turn derived from *MS2*$^C$ HTs but with a single C→U substitution in their loop that reverts them to Wt sequences described previously[48,49]. The sequences of all 1x and 2x *MS2* HT insertions used for SSA experiments are provided below, with color coding for flanking repeats and other features as follows:

*LYS2* repeats or *CAN1* repeats<br>
**TAA** or **TAG** (first in-frame stop codon)<br>
***MS2* HT** (with critical **Cyt/Thy** position)<br>
Protospacer<br>
**PAM**

1x *MS2*$^U$ + 1x *MS2*$^C$ (Upstream)<br>
CGAAGTTAAATCAAAAAATGACGATCCAAACTTTTTGAAAAAATTGAAAGATGTCATGCCTGCT<br>
GGTAAAGGTATGTTGAACGTTCAGCTACTAGTTGTT**TAA**AGAAAGA**ACATGAGGATTACCCATG**<br>
**T**CAGCAGCTCGACTGTAGAAA**ACATGAGGATCACCCATGT**AACGCTACCCATCCTGGTCGAGCT<br>
GGA**CGG**CGACTAGCGAAGTTAAATCAAAAAATGACGATCCAAACTTTTTGAAAAAATTGAAAGA<br>
TGTCATGCCTGCTGGTAAAGGTATGTTGAACGTTCAGCTACTAGTTGTT

1x *MS2*$^U$ + 1x *MS2*$^C$ (Downstream)<br>
CGAAGTTAAATCAAAAAATGACGATCCAAACTTTTTGAAAAAATTGAAAGATGTCATGCCTGCT<br>
GGTAAAGGTATGTTGAACGTTCAGCTACTAGTTGTT**TAA**CCCATCCTGGTCGAGCTGGA**CGG**CG<br>
ACAGAAAGA**ACATGAGGATTACCCATGT**CAGCAGCTCGACTGTAGAAA**ACATGAGGATCACCCA**<br>
**TGT**AACGCTATAGCGAAGTTAAATCAAAAAATGACGATCCAAACTTTTTGAAAAAATTGAAAGA<br>
TGTCATGCCTGCTGGTAAAGGTATGTTGAACGTTCAGCTACTAGTTGTT

2x *MS2*$^U$ (Upstream)<br>
CGAAGTTAAATCAAAAAATGACGATCCAAACTTTTTGAAAAAATTGAAAGATGTCATGCCTGCT<br>
GGTAAAGGTATGTTGAACGTTCAGCTACTAGTTGTT**TAA**AGAAAGA**ACATGAGGATTACCCATG**<br>
**T**CAGCAGCTCGACTGTAGAAA**ACATGAGGATTACCCATGT**AACGCTACCCATCCTGGTCGAGCT<br>
GGA**CGG**CGACTAGCGAAGTTAAATCAAAAAATGACGATCCAAACTTTTTGAAAAAATTGAAAGA<br>
TGTCATGCCTGCTGGTAAAGGTATGTTGAACGTTCAGCTACTAGTTGTT

2x *MS2*$^C$ (Upstream)<br>
CGAAGTTAAATCAAAAAATGACGATCCAAACTTTTTGAAAAAATTGAAAGATGTCATGCCTGCT<br>
GGTAAAGGTATGTTGAACGTTCAGCTACTAGTTGTT**TAA**AGAAAGA**ACATGAGGATCACCCATG**<br>
**T**CAGCAGCTCGACTGTAGAAA**ACATGAGGATCACCCATGT**AACGCTACCCATCCTGGTCGAGCT

GGA**CGG**CGACTAGCGAAGTTAAATCAAAAAATGACGATCCAAACTTTTTGAAAAAATTGAAAGA
TGTCATGCCTGCTGGTAAAGGTATGTTGAACGTTCAGCTACTAGTTGTT

1x *MS2*$^{U}$ (Upstream)

CGAAGTTAAATCAAAAAATGACGATCCAAACTTTTTGAAAAAATTGAAAGATGTCATGCCTGCT
GGTAAAGGTATGTTGAACGTTCAGCTACTAGTTGTT**TAA**AGAAGCTCGACTGTAGAAA**ACATGA**
**GGATTACCCATGT**AACGCTACCCATCCTGGTCGAGCTGGA**CGG**CGACTAGCGAAGTTAAATCAA
AAAATGACGATCCAAACTTTTTGAAAAAATTGAAAGATGTCATGCCTGCTGGTAAAGGTATGTT
GAACGTTCAGCTACTAGTTGTT

1x *MS2*$^{C}$ (Upstream)

CGAAGTTAAATCAAAAAATGACGATCCAAACTTTTTGAAAAAATTGAAAGATGTCATGCCTGCT
GGTAAAGGTATGTTGAACGTTCAGCTACTAGTTGTT**TAA**AGAAGCTCGACTGTAGAAA**ACATGA**
**GGATCACCCATGT**AACGCTACCCATCCTGGTCGAGCTGGA**CGG**CGACTAGCGAAGTTAAATCAA
AAAATGACGATCCAAACTTTTTGAAAAAATTGAAAGATGTCATGCCTGCTGGTAAAGGTATGTT
GAACGTTCAGCTACTAGTTGTT

2x *MS2*$^{U}$ (Upstream, *CAN1*)

ATATTCTGTCACGCAGTCCTTGGGTGAAATGGCTACATTCATCCCTGTTACATCCTCTTTCACA
GTTTTCTCACAAAGATTCCTTTCTCCAGCATTTGCT**TAG**AGAAAGA**ACATGAGGATTACCCATG**
**T**CAGCAGCTCGACTGTAGAAA**ACATGAGGATTACCCATGT**AACGCTACCCATCCTGGTCGAGCT
GGA**CGG**CGACTGAATATTCTGTCACGCAGTCCTTGGGTGAAATGGCTACATTCATCCCTGTTAC
ATCCTCTTTCACAGTTTTCTCACAAAGATTCCTTTCTCCAGCATTTG

Our *boxB* HTs were based off a 17 bp design that was previously employed for *boxB* arrays in eukaryotes[84,85]; these are identical to the 15 bp Wt *boxB* sequence within phage lambda's *nutR* element[121,122] except their stem region's complementarity is extended 1 bp with an additional pair of flanking G-C bases. For the 1x or 2x *boxB* designs, our 1x or 2x *MS2* designs listed above were modified by substituting the 19 bp *MS2* HTs with 17 bp *boxB* HTs. For the 3x *boxB* design, a third 17 bp *boxB* HT was added upstream of the protospacer-distal *boxB* with a 20 bp linker between them. We chose an A-rich sequence for this linker to reduce intramolecular base pairing potential. The sequences of all our *boxB* insertions in *LYS2* or the *EGFP* intron are provided below with color coding for features as follows:

*LYS2* repeat(s)
**TAA** (first in-frame stop codon)
***boxB* HT**
Protospacer
**PAM**
Flanking homology arms within *EGFP* (used for intron knock-in)

1x *boxB* (Upstream)

CGAAGTTAAATCAAAAAATGACGATCCAAACTTTTTGAAAAAATTGAAAGATGTCATGCCTGCT
GGTAAAGGTATGTTGAACGTTCAGCTACTAGTTGTT**TAA**AGAAGCTCGACTGTAGAA**GGCCCTG**
**AAAAAGGGCC**ACGCTACCCATCCTGGTCGAGCTGGA**CGG**CGACTAGCGAAGTTAAATCAAAAAA
TGACGATCCAAACTTTTTGAAAAAATTGAAAGATGTCATGCCTGCTGGTAAAGGTATGTTGAAC
GTTCAGCTACTAGTTGTT

2x *boxB* (Upstream)

CGAAGTTAAATCAAAAAATGACGATCCAAACTTTTTGAAAAAATTGAAAGATGTCATGCCTGCT
GGTAAAGGTATGTTGAACGTTCAGCTACTAGTTGTT**TAA**AGAAAG**GGCCCTGAAAAAGGGCC**AG
CAGCTCGACTGTAGAA**GGCCCTGAAAAAGGGCC**ACGCTACCCATCCTGGTCGAGCTGGA**CGG**CG
ACTAGCGAAGTTAAATCAAAAAATGACGATCCAAACTTTTTGAAAAAATTGAAAGATGTCATGC
CTGCTGGTAAAGGTATGTTGAACGTTCAGCTACTAGTTGTT

3x *boxB* (Upstream)

CGAAGTTAAATCAAAAAATGACGATCCAAACTTTTTGAAAAAATTGAAAGATGTCATGCCTGCT
GGTAAAGGTATGTTGAACGTTCAGCTACTAGTTGTT**TAA**AGAAAG**GGCCCTGAAAAAGGGCC**AA
CTAAACAACAAGAAAG**GGCCCTGAAAAAGGGCC**AGCAGCTCGACTGTAGAA**GGCCCTGAAAAAG**
**GGCC**ACGCTACCCATCCTGGTCGAGCTGGA**CGG**CGACTAGCGAAGTTAAATCAAAAAATGACGA
TCCAAACTTTTTGAAAAAATTGAAAGATGTCATGCCTGCTGGTAAAGGTATGTTGAACGTTCAG
CTACTAGTTGTT

1x *boxB* (Upstream, NAG PAM)

CGAAGTTAAATCAAAAAATGACGATCCAAACTTTTTGAAAAAATTGAAAGATGTCATGCCTGCT
GGTAAAGGTATGTTGAACGTTCAGCTACTAGTTGTT**TAA**AGAAGCTCGACTGTAGAA**GGCCCTG**
**AAAAAGGGCC**ACGCTACCCATCCTGGTCGAGCTGGA**CAG**CGACTAGCGAAGTTAAATCAAAAAA
TGACGATCCAAACTTTTTGAAAAAATTGAAAGATGTCATGCCTGCTGGTAAAGGTATGTTGAAC
GTTCAGCTACTAGTTGTT

1x *boxB* (Upstream, NGA PAM)

CGAAGTTAAATCAAAAAATGACGATCCAAACTTTTTGAAAAAATTGAAAGATGTCATGCCTGCT
GGTAAAGGTATGTTGAACGTTCAGCTACTAGTTGTT**TAA**AGAAGCTCGACTGTAGAA**GGCCCTG**
**AAAAAGGGCC**ACGCTACCCATCCTGGTCGAGCTGGA**CGA**CGACTAGCGAAGTTAAATCAAAAAA
TGACGATCCAAACTTTTTGAAAAAATTGAAAGATGTCATGCCTGCTGGTAAAGGTATGTTGAAC
GTTCAGCTACTAGTTGTT

1x *boxB* (Upstream, lacking downstream *LYS2* repeat)

CGAAGTTAAATCAAAAAATGACGATCCAAACTTTTTGAAAAAATTGAAAGATGTCATGCCTGCT
GGTAAAGGTATGTTGAACGTTCAGCTACTAGTTGTT**TAA**AGAAGCTCGACTGTAGAA**GGCCCTG**
**AAAAAGGGCC**ACGCTACCCATCCTGGTCGAGCTGGA**CGG**CGACTAG

1x *boxB* (Upstream, inverted protospacer)

CGAAGTTAAATCAAAAAATGACGATCCAAACTTTTTGAAAAAATTGAAAGATGTCATGCCTGCT
GGTAAAGGTATGTTGAACGTTCAGCTACTAGTTGTT**TAA**AGAAGCTCGACTGTAGAA**GGCCCTG**
**AAAAAGGGCC**ACGCTAAGTCG**CCG**TCCAGCTCGACCAGGATGGGAGCGAAGTTAAATCAAAAAA
TGACGATCCAAACTTTTTGAAAAAATTGAAAGATGTCATGCCTGCTGGTAAAGGTATGTTGAAC
GTTCAGCTACTAGTTGTT

3x *boxB* (Upstream, inverted protospacer)

CGAAGTTAAATCAAAAAATGACGATCCAAACTTTTTGAAAAAATTGAAAGATGTCATGCCTGCT
GGTAAAGGTATGTTGAACGTTCAGCTACTAGTTGTT**TAA**AGAAAG**GGCCCTGAAAAAGGGCC**AA
CTAAACAACAAGAAAG**GGCCCTGAAAAAGGGCC**AGCAGCTCGACTGTAGAA**GGCCCTGAAAAAG**
**GGCC**ACGCTAAGTCG**CCG**TCCAGCTCGACCAGGATGGGAGCGAAGTTAAATCAAAAAATGACGA
TCCAAACTTTTTGAAAAAATTGAAAGATGTCATGCCTGCTGGTAAAGGTATGTTGAACGTTCAG
CTACTAGTTGTT

2x *boxB* (Downstream)

CGAAGTTAAATCAAAAAATGACGATCCAAACTTTTTGAAAAAATTGAAAGATGTCATGCCTGCT
GGTAAAGGTATGTTGAACGTTCAGCTACTAGTTGTT**TAA**CCCATCCTGGTCGAGCTGGA**CGG**CG
ACAGAAAG**GGCCCTGAAAAAGGGCC**AGCAGCTCGACTGTAGAA**GGCCCTGAAAAAGGGCC**ACGC
TATAGCGAAGTTAAATCAAAAAATGACGATCCAAACTTTTTGAAAAAATTGAAAGATGTCATGC
CTGCTGGTAAAGGTATGTTGAACGTTCAGCTACTAGTTGTT

2x *boxB* (Upstream, embedded in *EGFP* intron V1.0)

GGGCGATGCCACCTACGGCAAGCTGACCCTGAAGTTCATCTGCACCACCGGGTAAGTACATACC
TCTCAGAACCCCCTCCGGCATGTATTCTTGTGTGCCTTGTCAACTCGTCAGACCGCGCAAATTT
CCGTCCGCAAAGAAAG**GGCCCTGAAAAAGGGCC**AGCAGCTCGACTGTAGAA**GGCCCTGAAAAAG**
**GGCC**ACGCTAAATCCGGTACGTGGCACTAACTAGTCTCTATTTTTCTTTTCATGAAGCTGCCCG
TGCCCTGGCCCACCCTCGTGACCACCC**TGA**CCTACGGCGT

For insertion of *boxB* HTs at the 3’ end of sgRNA scaffolds in paired-protospacer assays we modified a previously described sgRNA design with 3’ MS2$^{C}$ loops[123]. Its 2x 19 bp *MS2$^{C}$* HTs were replaced with 17 bp *boxB* HTs to generate our 2x *boxB* fusion. To facilitate synthesis of two or more nearby *boxB* HTs by IDT, the repetitiveness of our constructs was reduced with an A→G loop substitution in the second *boxB* HT that was previously found to maintain 89% of Wt activity in vivo[122]; in addition, the outermost G-C pair within the 17 bp HT was inverted to a C-G pair that maintains the putative stem length. To generate our 3x *boxB* fusion, both *MS2*$^{C}$ HTs were initially replaced with Wt *boxB* HTs, and then the *boxB** HT mutant was inserted between the first and third Wt *boxB* HTs at the second position along with 7 bp flanking linker sequences on each side. To generate our 1x *boxB* fusion, 34 bp that included the second HT were deleted from our 2x *boxB* construct, leaving 6 bp between the *SUP4* terminator and the remaining 17 bp Wt *boxB* HT. All of these designs retained additional base pairing potential beyond the 17 bp HT elements, which, might influence activity through a stem 'clamping' effect[124,125]. Note also that scaffold-internal HT insertions within the sgRNA were previously used in mammalian systems where 3' fusions failed[126]; 3' fusions such as ours might therefore require properties of the *SNR52* promoter, or other yeast properties, to function. Sequences are presented below with color coding for various features and spacer configurations:

*SNR52* promoter<br>
**Spacer 1**<br>
**Mismatches** relative to Protospacer 1<br>
**Spacer 2**<br>
sgRNA scaffold<br>
***boxB* or *boxB** HT** (**A->G** loop and **G-C -> C-G** stem substitutions)<br>
*SUP4* terminator

<u>3x *boxB* sgRNA fusion (0 Mm Spacer 1)</u>

TCTTTGAAAAGATAATGTATGATTATGCTTTCACTCATATTTATACAGAAACTTGATGTTTTCT<br>
TTCGAGTATATACAAGGTGATTACATGTACGTTTGAAGTACAACTCTAGATTTTGTAGTGCCCT<br>
CTTGGGCTAGCGGTAAAGGTGCGCATTTTTTCACACCCTACAATGTTCTGTTCAAAAGATTTTG<br>
GTCAAACGCTGTAGAAGTGAAAGTTGGTGCGCATGTTTCGGCGTTCGAAACTTCTCCGCAGTGA<br>
AAGATAAATGATC**CCCATCCTGGTCGAGCTGGA**GTTTTAGAGCTAGAAATAGCAAGTTAAAATA<br>
AGGCTAGTCCGTTATCAACTTGAAAAAGTGGCACCGAGTCGGTGCGGGAGC**GGCCCTGAAAAAG**<br>
**GGCC**GCCTACGAC**CGCCCTGAGAAAGGGCG**GTCGTAGCACGAGCG**GGCCCTGAAAAAGGGCC**CG<br>
CTCGTGTTCCCTTTTTTTGTTTTTTATGTCT

<u>3x *boxB* sgRNA fusion (5 Mm Spacer 1)</u>

TCTTTGAAAAGATAATGTATGATTATGCTTTCACTCATATTTATACAGAAACTTGATGTTTTCT<br>
TTCGAGTATATACAAGGTGATTACATGTACGTTTGAAGTACAACTCTAGATTTTGTAGTGCCCT<br>
CTTGGGCTAGCGGTAAAGGTGCGCATTTTTTCACACCCTACAATGTTCTGTTCAAAAGATTTTG<br>
GTCAAACGCTGTAGAAGTGAAAGTTGGTGCGCATGTTTCGGCGTTCGAAACTTCTCCGCAGTGA

AAGATAAATGATCAAACGCCTGGTCGAGCTGGAGTTTTAGAGCTAGAAATAGCAAGTTAAAATA
AGGCTAGTCCGTTATCAACTTGAAAAAGTGGCACCGAGTCGGTGCGGGAGCGGCCCTGAAAAAG
GGCCGCCTACGACCGCCCTGAGAAAGGGCGGTCGTAGCACGAGCGGGCCCTGAAAAAGGGCCCG
CTCGTGTTCCCTTTTTTTGTTTTTTATGTCT

3x *boxB* sgRNA fusion (20 Mm Spacer 1)

TCTTTGAAAAGATAATGTATGATTATGCTTTCACTCATATTTATACAGAAACTTGATGTTTTCT
TTCGAGTATATACAAGGTGATTACATGTACGTTTGAAGTACAACTCTAGATTTTGTAGTGCCCT
CTTGGGCTAGCGGTAAAGGTGCGCATTTTTTCACACCCTACAATGTTCTGTTCAAAAGATTTTG
GTCAAACGCTGTAGAAGTGAAAGTTGGTGCGCATGTTTCGGCGTTCGAAACTTCTCCGCAGTGA
AAGATAAATGATCAAACGAAGTTGATCTAGTTCGTTTTAGAGCTAGAAATAGCAAGTTAAAATA
AGGCTAGTCCGTTATCAACTTGAAAAAGTGGCACCGAGTCGGTGCGGGAGCGGCCCTGAAAAAG
GGCCGCCTACGACCGCCCTGAGAAAGGGCGGTCGTAGCACGAGCGGGCCCTGAAAAAGGGCCCG
CTCGTGTTCCCTTTTTTTGTTTTTTATGTCT

3x *boxB* sgRNA fusion (0 Mm Spacer 2)

TCTTTGAAAAGATAATGTATGATTATGCTTTCACTCATATTTATACAGAAACTTGATGTTTTCT
TTCGAGTATATACAAGGTGATTACATGTACGTTTGAAGTACAACTCTAGATTTTGTAGTGCCCT
CTTGGGCTAGCGGTAAAGGTGCGCATTTTTTCACACCCTACAATGTTCTGTTCAAAAGATTTTG
GTCAAACGCTGTAGAAGTGAAAGTTGGTGCGCATGTTTCGGCGTTCGAAACTTCTCCGCAGTGA
AAGATAAATGATCGTGCAGATGAACTTCAGGGTGTTTTAGAGCTAGAAATAGCAAGTTAAAATA
AGGCTAGTCCGTTATCAACTTGAAAAAGTGGCACCGAGTCGGTGCGGGAGCGGCCCTGAAAAAG
GGCCGCCTACGACCGCCCTGAGAAAGGGCGGTCGTAGCACGAGCGGGCCCTGAAAAAGGGCCCG
CTCGTGTTCCCTTTTTTTGTTTTTTATGTCT

1x *boxB* sgRNA fusion (0 Mm Spacer 1)

TCTTTGAAAAGATAATGTATGATTATGCTTTCACTCATATTTATACAGAAACTTGATGTTTTCT
TTCGAGTATATACAAGGTGATTACATGTACGTTTGAAGTACAACTCTAGATTTTGTAGTGCCCT
CTTGGGCTAGCGGTAAAGGTGCGCATTTTTTCACACCCTACAATGTTCTGTTCAAAAGATTTTG
GTCAAACGCTGTAGAAGTGAAAGTTGGTGCGCATGTTTCGGCGTTCGAAACTTCTCCGCAGTGA
AAGATAAATGATCCCCATCCTGGTCGAGCTGGAGTTTTAGAGCTAGAAATAGCAAGTTAAAATA
AGGCTAGTCCGTTATCAACTTGAAAAAGTGGCACCGAGTCGGTGCGGGAGCGGCCCTGAAAAAG
GGCCGCTCCCTTTTTTTGTTTTTTATGTCT

1x *boxB* sgRNA fusion (5 Mm Spacer 1)

TCTTTGAAAAGATAATGTATGATTATGCTTTCACTCATATTTATACAGAAACTTGATGTTTTCT
TTCGAGTATATACAAGGTGATTACATGTACGTTTGAAGTACAACTCTAGATTTTGTAGTGCCCT
CTTGGGCTAGCGGTAAAGGTGCGCATTTTTTCACACCCTACAATGTTCTGTTCAAAAGATTTTG
GTCAAACGCTGTAGAAGTGAAAGTTGGTGCGCATGTTTCGGCGTTCGAAACTTCTCCGCAGTGA
AAGATAAATGATCAAACGCCTGGTCGAGCTGGAGTTTTAGAGCTAGAAATAGCAAGTTAAAATA
AGGCTAGTCCGTTATCAACTTGAAAAAGTGGCACCGAGTCGGTGCGGGAGCGGCCCTGAAAAAG
GGCCGCTCCCTTTTTTTGTTTTTTATGTCT

1x *boxB* sgRNA fusion (0 Mm Spacer 2)

TCTTTGAAAAGATAATGTATGATTATGCTTTCACTCATATTTATACAGAAACTTGATGTTTTCT
TTCGAGTATATACAAGGTGATTACATGTACGTTTGAAGTACAACTCTAGATTTTGTAGTGCCCT
CTTGGGCTAGCGGTAAAGGTGCGCATTTTTTCACACCCTACAATGTTCTGTTCAAAAGATTTTG

GTCAAACGCTGTAGAAGTGAAAGTTGGTGCGCATGTTTCGGCGTTCGAAACTTCTCCGCAGTGA
AAGATAAATGATC**GTGCAGATGAACTTCAGGGT**GTTTTAGAGCTAGAAATAGCAAGTTAAAATA
AGGCTAGTCCGTTATCAACTTGAAAAAGTGGCACCGAGTCGGTGCGGGAGC**GGCCCTGAAAAAG**
**GGCC**GCTCCCTTTTTTTGTTTTTTATGTCT

2x *boxB* sgRNA fusion (0 Mm Spacer 1)

TCTTTGAAAAGATAATGTATGATTATGCTTTCACTCATATTTATACAGAAACTTGATGTTTTCT
TTCGAGTATATACAAGGTGATTACATGTACGTTTGAAGTACAACTCTAGATTTTGTAGTGCCCT
CTTGGGCTAGCGGTAAAGGTGCGCATTTTTTCACACCCTACAATGTTCTGTTCAAAAGATTTTG
GTCAAACGCTGTAGAAGTGAAAGTTGGTGCGCATGTTTCGGCGTTCGAAACTTCTCCGCAGTGA
AAGATAAATGATC**CCCATCCTGGTCGAGCTGGA**GTTTTAGAGCTAGAAATAGCAAGTTAAAATA
AGGCTAGTCCGTTATCAACTTGAAAAAGTGGCACCGAGTCGGTGCGGGAGC**GGCCCTGAAAAAG**
**GGCC**GCCACGAGCG**CGCCCTGAGAAAGGGCG**CGCTCGTGTTCCCTTTTTTTGTTTTTTATGTCT

2x *boxB* sgRNA fusion (5 Mm Spacer 1)

TCTTTGAAAAGATAATGTATGATTATGCTTTCACTCATATTTATACAGAAACTTGATGTTTTCT
TTCGAGTATATACAAGGTGATTACATGTACGTTTGAAGTACAACTCTAGATTTTGTAGTGCCCT
CTTGGGCTAGCGGTAAAGGTGCGCATTTTTTCACACCCTACAATGTTCTGTTCAAAAGATTTTG
GTCAAACGCTGTAGAAGTGAAAGTTGGTGCGCATGTTTCGGCGTTCGAAACTTCTCCGCAGTGA
AAGATAAATGATC**AAACGCCTGGTCGAGCTGGA**GTTTTAGAGCTAGAAATAGCAAGTTAAAATA
AGGCTAGTCCGTTATCAACTTGAAAAAGTGGCACCGAGTCGGTGCGGGAGC**GGCCCTGAAAAAG**
**GGCC**GCCACGAGCG**CGCCCTGAGAAAGGGCG**CGCTCGTGTTCCCTTTTTTTGTTTTTTATGTCT

2x *boxB* sgRNA fusion (0 Mm Spacer 2)

TCTTTGAAAAGATAATGTATGATTATGCTTTCACTCATATTTATACAGAAACTTGATGTTTTCT
TTCGAGTATATACAAGGTGATTACATGTACGTTTGAAGTACAACTCTAGATTTTGTAGTGCCCT
CTTGGGCTAGCGGTAAAGGTGCGCATTTTTTCACACCCTACAATGTTCTGTTCAAAAGATTTTG
GTCAAACGCTGTAGAAGTGAAAGTTGGTGCGCATGTTTCGGCGTTCGAAACTTCTCCGCAGTGA
AAGATAAATGATC**GTGCAGATGAACTTCAGGGT**GTTTTAGAGCTAGAAATAGCAAGTTAAAATA
AGGCTAGTCCGTTATCAACTTGAAAAAGTGGCACCGAGTCGGTGCGGGAGC**GGCCCTGAAAAAG**
**GGCC**GCCACGAGCG**CGCCCTGAGAAAGGGCG**CGCTCGTGTTCCCTTTTTTTGTTTTTTATGTCT

**Supplementary Note 3. Additional considerations for donor-dependent editing experiments in yeast.**

Cas9$^{NG}$ and Cas9$^{NG}$-2xN's phenotypic similarity in plating (Supplementary Fig. 6b) and liquid growth (Supplementary Fig. 7a–d) assays without lysine selection or donor DNA might reflect the sufficiency of a single unrepaired double-strand break to arrest cell division in yeast via the DNA damage response[127]. Under this interpretation, even if cleavage by native Cas9$^{NG}$ is delayed or infrequent relative to Cas9$^{NG}$-2xN under 1000nM conditions, it might reduce bulk growth to the same extent if both cleave at least once per generation in a similar number of cells. Addition of donor DNA during transformation partially alleviated their small-colony phenotypes under 1000nM plating conditions, that is, in some colonies but not others (Supplementary Fig. 6b), and we interpreted this as the result of occasional donor-mediated editing events that allow escape from subsequent re-cleavage. The above interpretations assume that our NG variants, unlike Wt variants, do not outright suppress colony formation by unedited cells because they do not cleave actively enough, i.e., they cannot cleave in ~ 100% of plated cells.

When we quantified the initial transformation-associated editing fractions from donor-treated samples by plating in parallel to select for *LYS2*$^{+}$ colonies immediately, we could not detect significant differences between Cas9$^{NG}$ and Cas9$^{NG}$-2xN at either 15nM or 1000nM β-E (Supplementary Fig. 6c). Anticipating this possibility, we grew the same transformant populations in parallel without lysine selection on separate plates containing 0, 15, or 1000nM β-E, and then scrape-homogenized the colonies after 44 h growth for replating to quantify *LYS2*$^{+}$ fractions. Small but significant increases in editing were detected for $Cas9^{NG}$-2xN at 0nM and 15nM β-E in the presence of donor DNA (Supplementary Fig. 6d). No such advantage was detected with Cas9-2xN when the same transformations were grown at 1000nM β-E before replating: virtually 100% of the viable population was *LYS2*$^{+}$ in both native $Cas9^{NG}$ and $Cas9^{NG}$-2xN populations, as measured for $Cas9^{Wt}$-2xN under these conditions (Supplementary Fig. 6e). Control transformations carried out under identical conditions of 1000nM outgrowth confirmed that editing requires both donor DNA and a targeting sgRNA, as the absence of either yielded zero *LYS2*$^{+}$ colonies upon replating (Supplementary Fig. 6e). Evidently, the similar growth phenotypes suffered by unedited parental cells with either $Cas9^{NG}$ or $Cas9^{NG}$-2xN at 1000nM β-E allowed edited cells which escape cleavage to comparably overtake their population on plates, though the edited fractions from those transformations were initially < 20% (Supplementary Fig. 6c). Although we did not detect bulk growth defects with 0nM and 15nM β-E for either $Cas9^{NG}$ or $Cas9^{NG}$-2xN in the absence of lysine selection, it is still formally possible that sub-detectable targeting activity during outgrowth depleted unedited cells to some extent, and perhaps more so with the $Cas9^{NG}$-2xN fusion. In that scenario, $Cas9^{NG}$-2xN's apparent increases in editing after replating from 0nM and 15nM β-E scrapes could have arisen from a slight advantage at enriching for edited cells during outgrowth, at least in part.

**Supplementary Note 4. Additional considerations for interpreting the effects of protospacer orientation and positioning relative to *cis*-interacting elements.**

Despite expectations that our protospacer with the PAM-downstream orientation would be prone to RNAP-mediated dislodging[73] and license editing at higher levels than its inverted (PAM-upstream) counterpart (Supplementary Fig. 8a), our tests with the weak *pLYS2* promoter revealed comparable editing with native Cas9$^{MT3}$ and Cas9$^{NG}$ in both protospacer orientations (Supplementary Fig. 8b). One plausible interpretation is that recruitment occurs slower than post-cleavage dislodging for native Cas9$^{MT3}$ and Cas9$^{NG}$ under these conditions and is thus rate-limiting for them, such that they only rarely rebind if dislodged, while the reverse is true for Cas9$^{MT3}$-2xN and Cas9$^{NG}$-2xN: higher recruitment rates stimulated by TraCT allow for additional cycles of binding, cleavage, and dislodging until the target is edited, but not when dislodging is inherently rare (as with PAM-upstream protospacers). It was not determined whether attenuated sgRNAs [128], such as mismatched sgRNAs that are dislodged by RNAPs even in the PAM-upstream orientation[129], can license TraCT where standard sgRNAs do not. Note that other cellular factors, including the FACT complex[130] found in mammals and yeast[131], could contribute to basal Cas9 dislodging and thus editing frequencies in our assays. In addition, we tested *pLYS2* with the PAM-downstream orientation and two downstream *boxB* HTs and found a small but significant increase for Cas9$^{MT3}$-2xN (Supplementary Fig. 8b); therefore, positioning the PAM between its protospacer and HTs does not preclude transcription-associated editing either. Nevertheless, it should be noted that our SSA editing experiments do not formally distinguish between facilitated Cas9 recruitment at the PAM-recognition step and facilitated Cas9 dislodging post-cleavage, given that in vivo repair outcomes may be biochemically rate-limited by either. Furthermore, it remains to be determined whether TraCT could be adapted for use with the Cas9 ortholog from *Staphylococcus aureus* (SauCas9) that was reported to display inherent multiple-turnover cleavage activity in vitro[132], and whether protospacer orientation would influence editing in that scenario.

**Supplementary Note 5. Additional considerations for SSA experiments with Cas9(-scTetR) and *trans*-expressed TetR.**

The stimulatory effect of *trans*-expressed TetR on native $Cas9^{MT3}$ activity was greatest with moderate or high levels of TetR and in all cases was significantly reduced upon Dox addition, roughly to baseline levels (Supplementary Fig. 9e). TetR might locally increase access to naked DNA by displacing nucleosomes, as seen for Reb1 binding sites in yeast[90] and as proposed for the proxy-CRISPR effect[92]. Conversely, we noticed that maximal $Cas9^{MT3}$-scTetR activity in the absence of Dox was slightly reduced by *trans* TetR expression, particularly at moderate and high levels, though no such effects were seen with $Cas9^{Wt}$-scTetR (Supplementary Figs. 9e–f). This apparent inhibitory effect could arise if concentrations of TetR are high enough to outcompete $Cas9^{MT3}$-scTetR for binding at *tetO2*. In other words, while the covalent linkage used in our $Cas9^{MT3}$-scTetR fusions may provide a superior recruitment advantage over the putative nucleosome displacement effects afforded to native $Cas9^{MT3}$ by TetR alone, this advantage could be masked if moderate and high levels of TetR outcompete scTetR for *tetO2* binding. Consistent with the notion of a recruitment advantage for $Cas9^{MT3}$-scTetR, covalently linked Cas9-Cas9 fusions were previously reported to outperform independently expressed Cas9 proteins[113].

**Supplementary Note 6. Additional considerations for control data from paired-protospacer experiments examining the effects of anchored boxB hairpins.**

In control experiments where boxB hairpins were absent from both sgRNAs, 2xN fusions did not increase editing (Fig. 6b). The basal activity of native Cas9$^{Wt}$ with or without sgRNA-boxB fusions, as well as Cas9$^{Wt}$-2xN with sgRNAs that lacked boxB, was notably somewhat higher with 5 Mm in spacer 1 relative to 20 Mm (Fig. 6b). We attribute this to the proxy-CRISPR effect described previously[92]. Importantly, only background editing was detected when spacer 2 was effectively non-targeting with 15 Mm and spacer 1 contained 5 Mm; this confirmed that the latter 5 Mm are sufficient to abolish cleavage activity at protospacer 1 (Fig. 6b).

We also probed whether the anchored hairpin-stimulated activity of Cas9-2xN at paired-protospacer targets may require particular protospacer orientations (Fig. 6a; Supplementary Fig. 12a), reasoning that static positioning of the spacer 1 R-loop on DNA would produce similar constraints as those limiting scTetR with certain *tetO2* configurations (Supplementary Fig. 8c–d). Indeed, when protospacer 2 was inverted without altering protospacer 1 or the distance between them (Supplementary Fig. 12e), the 2xN fusion had no discernible editing advantage with three boxB hairpins in either sgRNA configuration (Supplementary Fig. 12f). Paired targeting with Cas9-Cas9 fusions was also previously found to depend on certain protospacer orientations[113], though the molecular basis for this remains poorly understood. These data argue that facilitated recruitment models which only consider the effect of increasing Cas9’s local concentration near target DNA are insufficient to explain how subPAM interactions are functionally complemented in eukaryotes. Rather, such models might also consider concomitant nucleosome displacement effects that temporarily reduce DNA occlusion (see Discussion).
